# Domain-resolved epitope mapping of B7-H3-targeting VHH-Fcs defines isoform-selective binders for antibody–drug conjugate development

**DOI:** 10.64898/2026.10.16.759583

**Authors:** Xiang Fang, Benjamin Chalopin, S. Seyda Gokturk, Ankita S. Bhagwani, Milana Sudareva, Ronja Molkenbur, Wenjing Tian, Min Feng, Fenna L.H. van Rijt, Kim van Noort, Jannet Koelewijn, Marianthi Chatziandreou, Baojie Zhang, Johannes H. M. Merks, Claudia Rossig, Ramona Meissner, Claudia Y. Janda

## Abstract

The immune checkpoint molecule B7-H3 has emerged as one of the most actively pursued therapeutic targets in solid tumors, and multiple B7-H3-targeting antibody–drug conjugates have entered clinical evaluation. B7-H3 occurs as a 2Ig isoform carrying a single IgV-IgC domain pair and a predominant 4Ig isoform carrying two tandem pairs of near-identical sequence in their extracellular domains. Resolving which pair an antibody engages, or how antibodies distinguish the two isoforms, requires epitope mapping at residue level. Here we generated a panel of six B7-H3-targeting VHH-Fcs and characterized them by surface plasmon resonance, epitope binning, domain-resolved deep mutational scanning and receptor internalization, benchmarked against the clinically advanced antibodies ifinatamab, mirzotamab and vobramitamab. Most VHH-Fcs bound the membrane-proximal pair more tightly than the distal pair, and 4Ig 10-to 100-fold more tightly than 2Ig, whereas the three clinical antibodies showed no such discrimination. Displaying each domain pair independently on yeast identified the residues that account for this selectivity. Internalization varied several-fold among binders, including between VHH-Fc 620 and 636, which have near-identical footprints and comparable affinities. VHH-Fc 620 internalized most efficiently, also compared to ifinatamab, and was conjugated to exatecan through a protease-cleavable linker. The conjugate showed B7-H3-dependent cytotoxicity that scaled with B7-H3 cell surface density, though it was less potent than ifinatamab-exatecan. Affinity, epitope and internalization therefore behaved as independent characteristics rather than integrating to predict antibody–drug conjugate potency. Together, these results define the molecular basis for isoform-selective recognition of B7-H3 and identify VHH-Fc 620 as a starting point for isoform-selective targeting.

## Introduction

B7-H3 (CD276), a member of the B7 family of immune regulatory proteins, has emerged as an attractive target for cancer therapy owing to its widespread expression across a broad range of solid tumors and its relatively restricted expression in normal tissues. Elevated B7-H3 expression has been reported in over 60 % and up to 93 % of patient tumor tissues in the vast majority of cancer types^1^. Although initially described as an immune checkpoint molecule, the physiological receptor(s) and biological function of B7-H3 remain incompletely understood. Despite this uncertainty, accumulating evidence suggests that B7-H3 contributes to tumor progression by promoting proliferation, invasion, angiogenesis, metabolic adaptation, and resistance to therapy^2,3^.

At the molecular level, human B7-H3 (hB7-H3) exists as two isoforms generated by alternative splicing. The 2Ig isoform contains a single IgV–IgC domain pair in its extracellular domain (ECD), and the predominant 4Ig isoform comprises two tandem IgV–IgC domain pairs^4,5^. In addition to these membrane-bound isoforms, soluble B7-H3 has also been described, arising through either alternative splicing or proteolytic shedding^6,7^. The alternatively spliced soluble form corresponds to the 2Ig ectodomain, with four additional residues at its C-terminus^6^. Isoform-selective targeting has been proposed as a way to limit sequestration by circulating soluble 2Ig, and MIL33B, an antibody with 8-fold selectivity for 4Ig over 2Ig, has recently been developed for radioligand therapy^8^. VHHs against B7-H3 have also been generated for CAR T-cell applications, though these engage both IgV-IgC domain pairs and both isoforms^5^. In both cases, the molecular basis for isoform discrimination, and whether binders can distinguish the two near-identical domain pairs within 4Ig, remain undefined.

B7-H3 is now pursued across multiple therapeutic modalities, including unconjugated monoclonal antibodies (mAbs), antibody–drug conjugates (ADCs), bispecific antibodies and chimeric antigen receptor (CAR) T-cell therapies, several of which have entered clinical evaluation^9^. Among these approaches, ADCs have shown particularly encouraging clinical progress. The most advanced candidate, ifinatamab deruxtecan, has been evaluated in Phase III clinical trials for small-cell lung cancer (NCT06203210, initiated in 2024) and metastatic castration-resistant prostate cancer (NCT06925737, initiated in 2025). Another clinically advanced ADC, vobramitamab duocarmazine, has been evaluated in Phase II studies across multiple solid tumor indications, including prostate cancer, head and neck squamous cell carcinoma, and small cell lung cancer (NCT05551117, initiated in 2023).

Despite the rapid expansion of B7-H3-targeted therapies, little is known about how the molecular properties of anti-B7-H3 antibodies determine their suitability for different therapeutic applications. In contrast to well-characterized targets such as HER2 or EGFR, for which extensive structure-function relationships have been established, B7-H3 antibodies have undergone relatively limited comparative characterization^10,11^. Affinity influences biodistribution and retention at the target site^12^, while mAbs recognizing distinct epitopes on the same target can differ in their ability to trigger receptor internalization and intracellular trafficking toward lysosomal compartments, where most cleavable-linker ADCs release their payload^13^. Differences in receptor clustering, endocytic routing, or recycling may therefore affect intracellular payload delivery^14^. Furthermore, ADC efficacy depends on factors extending beyond the antibody itself, including antigen density, internalization kinetics, intracellular trafficking, linker chemistry, payload mechanism of action, and tumor-specific characteristics^15–17^. These characteristics are interdependent, and individually they are poor predictors of ADC performance, which is why comparative, target-specific characterization of mAb panels is needed.

Here, we generated a panel of B7-H3-targeting VHH-Fcs and evaluated their internalization ability to predict their suitability for ADC development. VHHs are attractive for targeted drug delivery, combining high stability and small size with a modular architecture that supports engineering into multivalent and multifunctional formats^18^. We characterized their affinity, epitope footprint and receptor-mediated internalization properties, and benchmarked them against three clinically advanced B7-H3-targeting mAbs, ifinatamab, mirzotamab and vobramitamab. Because the two IgV-IgC domain pairs of 4Ig hB7-H3 are highly similar in sequence, conventional epitope analysis cannot resolve which pair a binder engages. We therefore combined surface plasmon resonance (SPR)-based epitope binning with domain-resolved deep mutational scanning (DMS), displaying each domain pair independently on yeast to map binding at the level of individual residues. VHH-Fc 620 was then conjugated to exatecan through a protease-cleavable linker and compared with an ifinatamab-based ADC bearing the same linker and payload across cell lines expressing different levels of B7-H3. Together, this work identifies B7-H3 binders that discriminate between the two domain pairs of 4Ig and between 4Ig and 2Ig, defines the residues underlying this selectivity, and establishes VHH-Fc 620 as a candidate for further ADC development.

## Results

### Isolation and characterization of hB7-H3-targeting VHHs

To generate anti-human B7-H3 VHHs, two independent llama immunization campaigns were performed (*n*=2 animals per campaign) using the recombinant ECD of B7-H3, administered by subcutaneous injection (Figure 1a). Immune VHH libraries were constructed from peripheral blood lymphocytes of the immunized animals and subjected to two consecutive rounds of phage display panning on 4Ig hB7-H3 ECD. Positive clones were identified by phage ELISA and sequenced. Selected VHHs were reformatted into a bivalent VHH-Fc format and produced by transient expression in Expi293F cells followed by Protein A affinity purification and size-exclusion chromatography (SEC). Through iterative rounds of sequence analysis, expression, and biochemical characterization, six candidates (VHH-Fc 620, 621, 622, 628, 633, and 636) were prioritized on the basis of CDR sequence diversity, purification yield, purity by SDS-PAGE, monodispersity by analytical SEC, and thermal stability (Supplementary Figure S1a-b). By ELISA, all six bound plate-immobilized 4Ig hB7-H3 in a concentration-dependent manner, with apparent EC50 values for the bivalent VHH-Fc constructs in the subnanomolar range (0.15-0.69 nM; Figure 1b).

**Figure 1.**
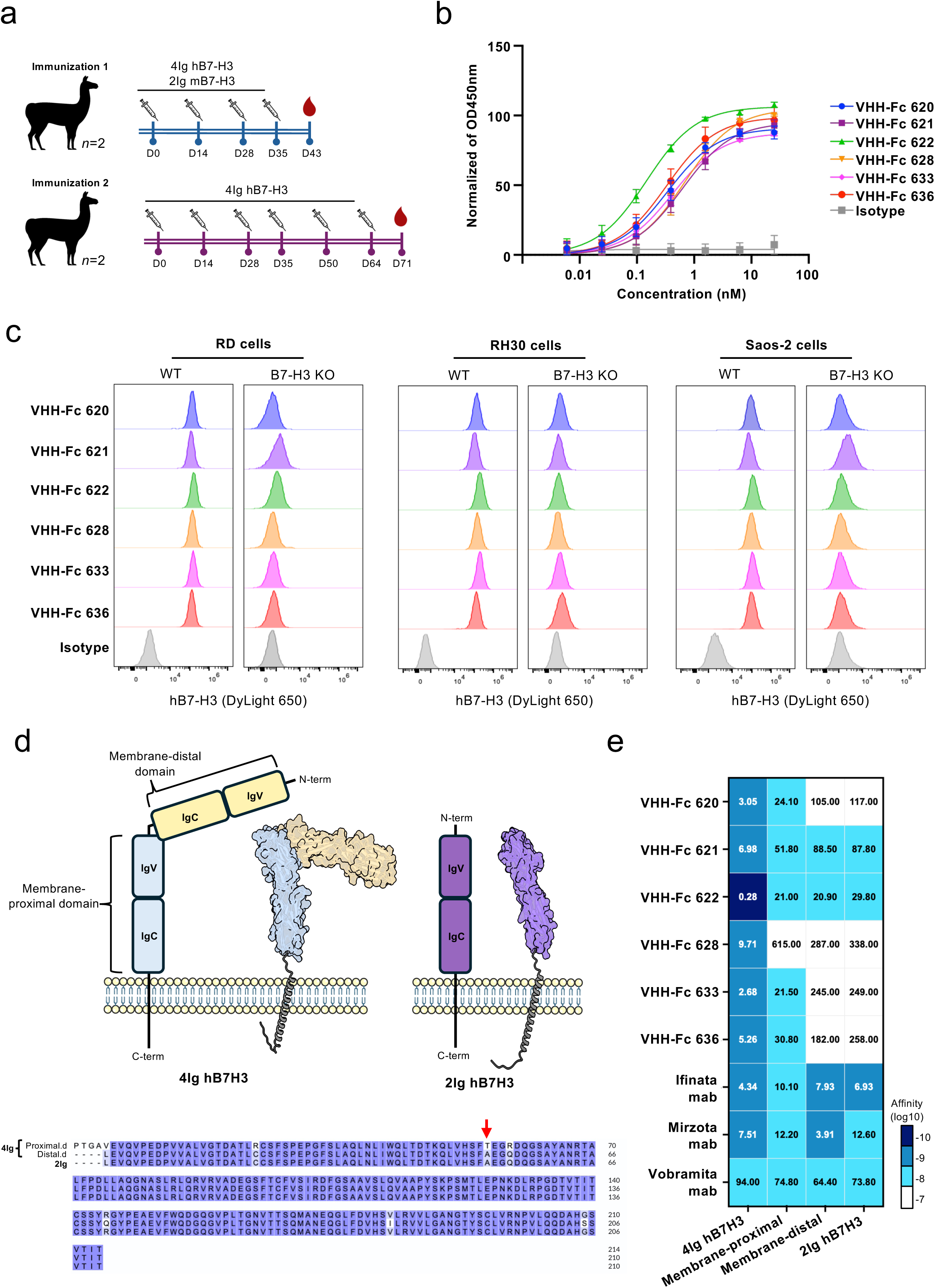
Isolation and characterization of human B7-H3-targeting VHHs. **(a)** Immunization scheme. Two llamas per campaign received subcutaneous injections of recombinant B7-H3 ECD (100 µg per antigen for the first two injections, 50 µg thereafter). Immunization 1: four injections of 4Ig human B7-H3 (hB7-H3) co-administered with 2Ig mouse B7-H3 (mB7-H3). Immunization 2: six injections of 4Ig hB7-H3. Immune libraries from both campaigns were subjected to phage display selection, and the resulting clones were screened by phage ELISA. **(b)** Binding of selected VHH-Fcs to plate-immobilized recombinant 4Ig hB7-H3, determined by ELISA. OD450 values were normalized to signal at 25 nM. Data are mean ± SD of three independent experiments, each performed in technical triplicate. **(c)** Binding of selected VHH-Fcs (40 nM) to wild-type (WT) and B7-H3-knockout (KO) RD, RH30, and Saos-2 cells determined by flow cytometry. Bound VHH-Fc was detected with a DyLight 650-conjugated anti-human IgG secondary antibody. Data shown are from one representative experiment of three independent experiments with similar results. **(d)** AlphaFold-predicted structures of the 4Ig and 2Ig hB7-H3 isoforms, shown alongside a schematic of their domain organization. Below, sequence alignment of the membrane-proximal and membrane-distal IgV-IgC domain pair of 4Ig and 2Ig hB7-H3 (>96% sequence identity). Amino acid T298 in the membrane proximal domain of 4Ig hB7-H3 and its equivalent position in the membrane-distal domain of 4Ig and 2Ig is indicated with an arrow. **(e)** Heatmap of apparent binding affinities (KD) for the VHH-Fcs and three benchmarking mAbs against 4Ig hB7-H3, its isolated membrane-proximal and membrane-distal domains, and 2Ig hB7-H3, determined by SPR (corresponding sensograms are shown in Supplementary Figure S3). Values within cells are KD in nM; the color scale indicates log10 (KD in M). Data are the mean of three independent experiments with similar results.

We next evaluated the binding of these VHH-Fcs toward membrane-expressed hB7-H3 using three human sarcoma cell lines (RH30, RD, and Saos-2), all known to express hB7-H3^5,19^. For each line, we generated a corresponding hB7-H3 knockout (KO) line to rigorously assess specificity (Supplementary Figure S2a-c). Flow cytometry demonstrated that all VHH-Fcs bound the hB7-H3 wild-type (WT) cells, with no detectable staining of the KO cells above isotype control, confirming that all six bind membrane-expressed hB7-H3 and that binding is B7-H3 specific (Figure 1c).

To characterize these interactions in more detail, we determined the binding kinetics of the VHH-Fcs toward the 4Ig and 2Ig hB7-H3 isoforms by SPR (Supplementary Figure S3, Supplementary Table S1), and included ifinatamab, mirzotamab, and vobramitamab, three clinically advanced B7-H3-targeting mAbs^20–22^, for benchmarking (Figure 1d-e). Because the two IgV-IgC domain pairs of 4Ig, designated membrane-proximal and membrane-distal domains, share high sequence identity, we also evaluated binding of the VHH-Fcs toward the individual IgV-IgC domains.

SPR analysis demonstrated that VHH-Fc 620, 621, 628, 633, and 636 bound 4Ig hB7-H3 with single-digit nanomolar affinities, whereas VHH-Fc 622 exhibited subnanomolar affinity, driven primarily by rapid association and dissociation. Ifinatamab and mirzotamab bound in the same single-digit nanomolar range, while vobramitamab bound 4Ig hB7-H3 with a KD of 94.0 nM, approximately 10-to 30-fold more weakly than most of the panel.

Analysis of the isolated domains revealed differential domain recognition within the VHH-Fc panel and the benchmarking mAbs. Four of the six VHH-Fcs preferentially bound the membrane-proximal domain. VHH-Fc 633, 636, and 620 bound the membrane-proximal domain with double-digit nanomolar affinity but the membrane-distal domain 4-to 12-fold more weakly, in the triple-digit nanomolar range. VHH-Fc 621 showed the same preference more modestly (approximately 2-fold). In contrast, VHH-Fc 622 and 628 displayed comparable affinities toward both domains. VHH-Fc 622 retained double-digit nanomolar affinity for both domains, and VHH-Fc 628 bound both domains weakly. The benchmarking mAbs ifinatamab, mirzotamab, and vobramitamab bound both isolated domains with similar affinities, consistent with those measured against 4Ig hB7-H3, indicating no comparable domain discrimination. For both panels, affinities for 2Ig closely tracked those for the isolated membrane-distal domain. Most VHH-Fcs bound 2Ig 10-to 100-fold more weakly than 4Ig, whereas the benchmarking mAbs bound the two isoforms with comparable affinities.

Together, these data identify six B7-H3-specific VHH-Fcs. Four VHH-Fcs, unlike the clinical benchmarking mAbs, largely discriminate between the two IgV-IgC domain pairs of 4Ig and between 4Ig and 2Ig. As this discrimination may limit sequestration of the VHH-Fcs by soluble 2Ig hB7-H3, we set out to define the epitopes of the panel at higher resolution.

### SPR-based epitope binning identifies three distinct epitope bins

Because this discrimination implies that the VHH-Fcs engage surface residues on hB7-H3 that differ between the membrane-distal and membrane-proximal IgV-IgC domains and between 4Ig and 2Ig, we first used SPR-based epitope binning to establish how the VHH-Fc binding epitopes are distributed across the ECD surface relative to those of the benchmarking mAbs. We performed pairwise binding experiments for all VHH-Fc and mAb combinations using a capture-based assay format (Figure 2a). The membrane-proximal IgV–IgC domain was selected as the antigen so that each binder could engage only one copy of its epitope, avoiding the avidity and steric effects that arise from the duplicated 4Ig architecture that would otherwise confound the assignment of epitope bins. In addition, most VHH-Fcs bound this domain more strongly (Figure 1e), giving stronger signals for binning. Representative sensorgrams illustrating antigen capture and subsequent binding of the second binder are shown in Supplementary Figure S4. Binder pairs capable of simultaneous binding generated an increase in response signal upon injection of the second binder, whereas competing pairs showed little or no signal enhancement, consistent with recognition of sterically overlapping or spatially adjacent epitopes (Figure 2a-b).

**Figure 2.**
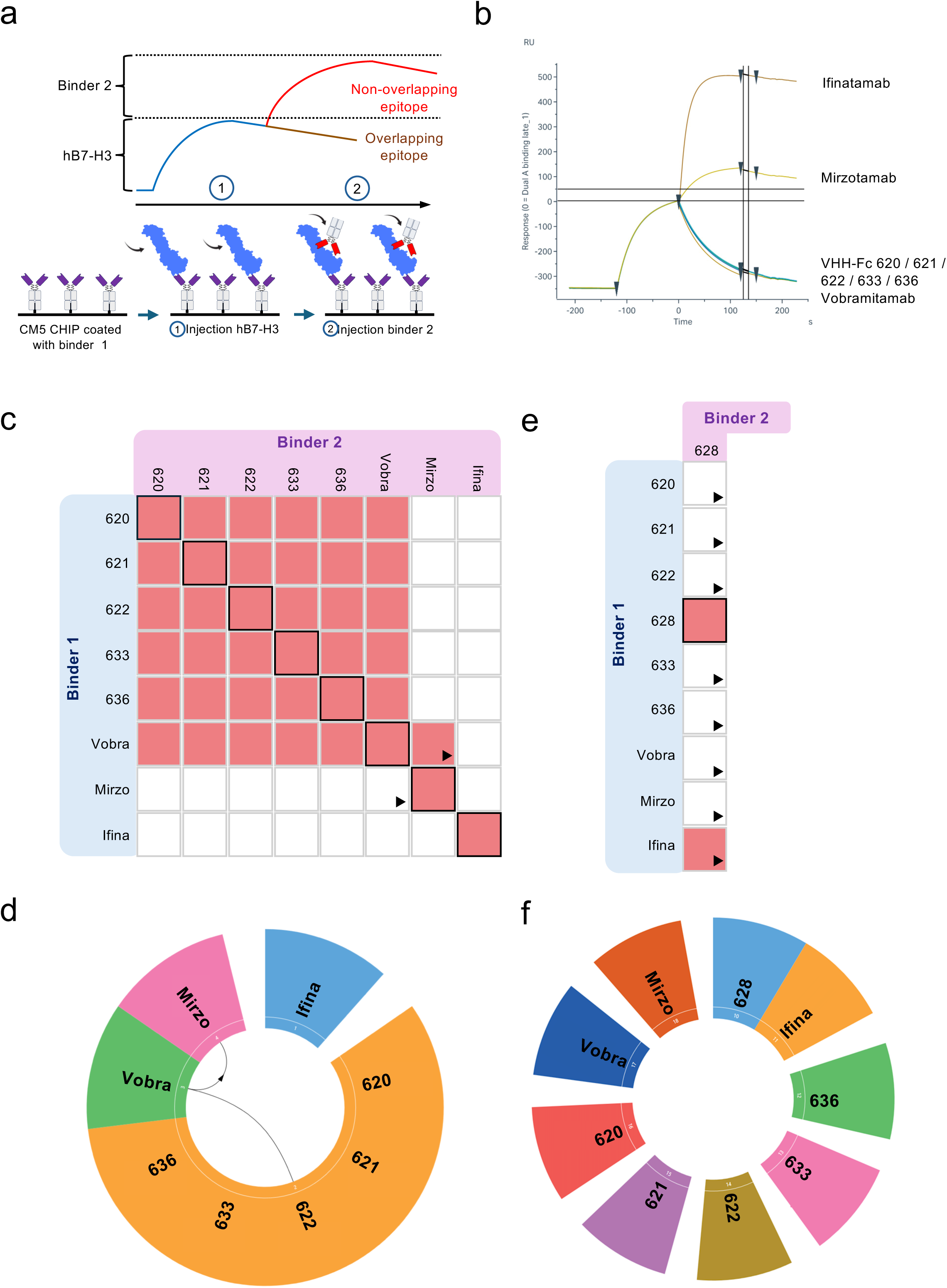
SPR-based epitope binning of B7-H3-binding VHH-Fcs and benchmarking mAbs. **(a)** Schematic overview of SPR-based epitope binning. The first binder is immobilized on a CM5 sensor chip via amine coupling. The full-length or membrane-proximal domain of 4Ig hB7-H3 is subsequently injected, and binding is monitored. The second binder is then injected, and changes in binding are recorded. Non-overlapping epitopes result in additional binding, whereas overlapping epitopes lead to competitive blocking, preventing further binding. **(b)** Example of data acquisition for SPR-based epitope binning. VHH-Fc 620 is immobilized on a CM5 sensor chip. Upon injection of the membrane-proximal domain of 4Ig, an increase in binding signal is observed. The second binder is then injected. Binders recognizing the same or overlapping epitopes do not produce an additional increase in signal (e.g., VHH-Fc 620, 621, 622, 633, 636, and mirzo/mirzotamab), whereas binders targeting distinct epitopes result in a further increase in binding (e.g., ifina/ifinatamab, vobra/vobramitamab). **(c)** Pairwise competition matrix for the VHH-Fcs (excluding VHH-Fc 628) and mAbs, using the membrane-proximal domain of 4Ig hB7-H3 as antigen. Red squares indicate competition (shared or overlapping epitopes); white squares indicate no competition (distinct epitopes). Red squares with black borders are self-competition controls. Arrows depict representative competitive interactions between binders in different bins. **(d)** Wheel visualization of the binning analysis in (c). Colored sectors correspond to epitope bins, with labels indicating the binders in each bin. **(e)** Pairwise competition matrix for VHH-Fc 628, using full-length 4Ig hB7-H3 as antigen, as VHH-Fc 628 bound the isolated membrane-proximal domain too weakly for reliable binning. **(f)** Wheel visualization of the binning analysis in (e).

Epitope binning segregated the nine binders into three distinct epitope bins (Figure 2c-f). Five VHH-Fcs (620, 621, 622, 633, and 636) clustered within bin 1, together with vobramitamab. Mirzotamab occupied a second bin positioned adjacent to that of vobramitamab. Notably, vobramitamab prevented subsequent binding of mirzotamab, whereas mirzotamab did not interfere with vobramitamab binding. This asymmetric competition pattern suggests that the two mAbs recognize neighboring epitopes and that binding of vobramitamab may sterically restrict access of mirzotamab to its binding site.

VHH-Fc 628 could not be binned on the isolated membrane-proximal domain owing to its low affinity and rapid dissociation rate, so binning was performed on full-length 4Ig hB7-H3. Avidity from the duplicated domains of 4Ig substantially reduced the apparent dissociation rate and enabled robust bin assignment (Figure 2e-f, Supplementary Figure S5). On this antigen, VHH-Fc 628 segregated into a third bin, shared with ifinatamab. Together, these data showed that the VHH-Fc panel occupies two epitope regions of hB7-H3, each also engaged by a clinically advanced mAb.

### DMS defines the molecular determinants of hB7-H3 recognition

While epitope binning groups binders by competition, it does not identify the involved contact residues. To resolve the epitopes at residue level, and to determine which residues underlie the domain and isoform discrimination observed by SPR, we performed DMS coupled with yeast surface display of the membrane-proximal and membrane-distal domains of 4Ig hB7-H3. The domains were displayed independently to enable domain-resolved epitope mapping, as previously described^23^ (Supplementary Figure S6a-d). VHH-Fc binding affinities to yeast-displayed IgV-IgC domains were validated against the SPR data, confirming consistent domain-specific binding profiles (Supplementary Figure S7a-e, Supplementary Table S2). Since all VHH-Fcs except 628 displayed equal or higher affinity for the membrane-proximal domain, epitope mapping was performed on this domain for the entire panel (see Materials and Methods, Supplementary Figure S8a-b). Mutation-sensitive residues were then mapped onto an AlphaFold3 model of the IgV-IgC domain to visualize the VHH-Fc footprints. While most residues were surface exposed, a subset was located within the domain core, suggesting that their substitution impairs local folding or conformational stability rather than directly disrupting the VHH-binding interface.

Mapping of the VHH-Fcs showed that five of the six (VHH-Fc 620, 621, 622, 633 and 636) recognized overlapping epitopes within the IgV-like domain, mostly involving β-strand residues (Figure 3a-b, Supplementary Figure S9). VHH-Fc 622 targeted the same general region but exhibited a more restricted pattern of mutation sensitivity, consistent with its faster binding kinetics. In contrast, VHH-Fc 628 recognized a distinct epitope in the central region of the IgC-like domain.

**Figure 3.**
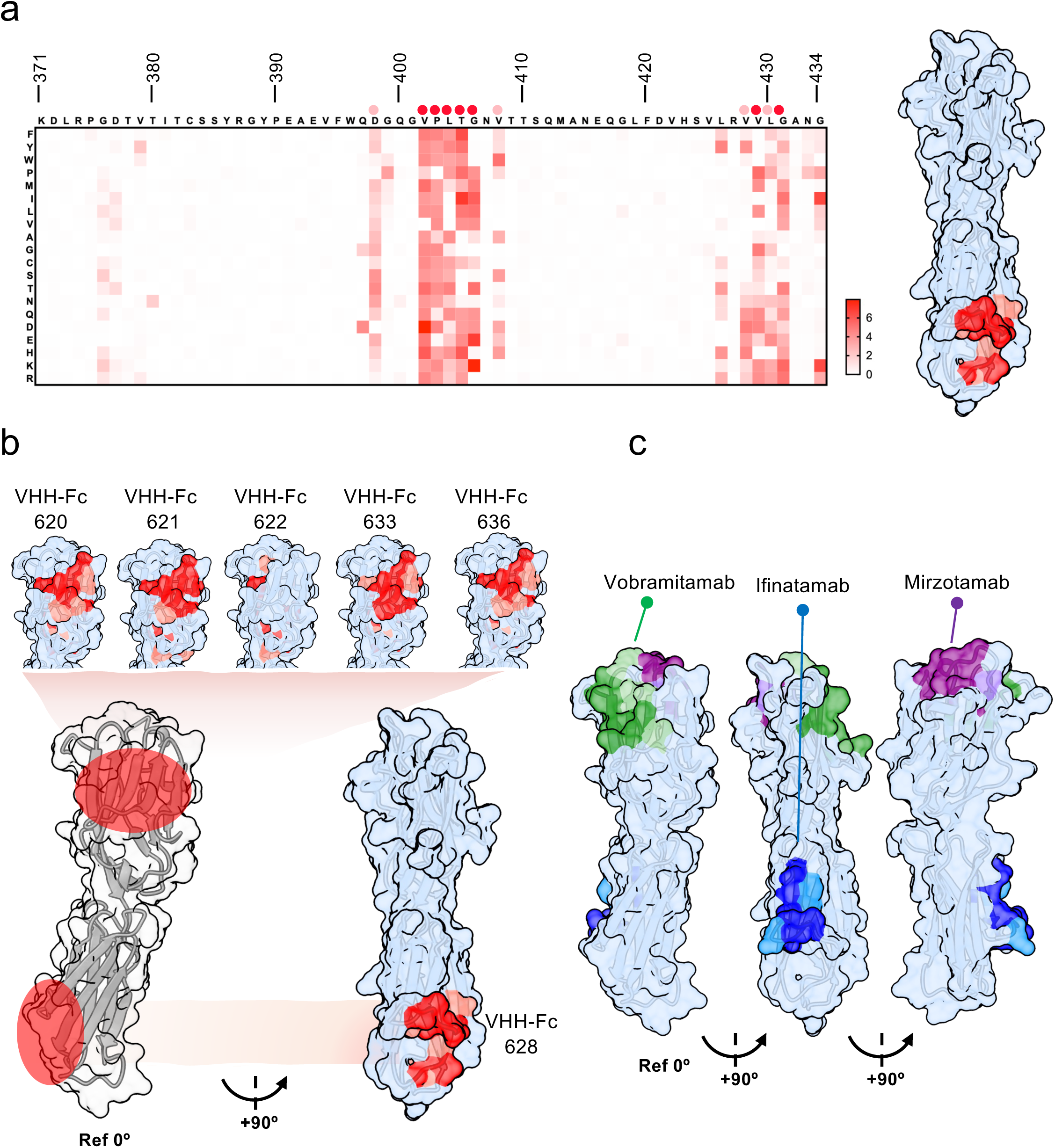
Functional epitopes of VHH-Fcs and benchmarking mAbs on hB7-H3. **(a)** Heatmap showing the effect of single amino acid substitutions within the IgC-like region of the hB7-H3 membrane-proximal domain (residues 371–434; UniProt Q5ZPR3, CD276_HUMAN) on VHH-Fc 628 binding. Critical residues are highlighted on the structural representation of the hB7-H3 membrane-proximal domain: dark red denotes residues for which ≥50% of substitutions tested (n = 19) disrupted binding, whereas light red denotes residues for which ≥25% disrupted binding. **(b)** Surface representations of the hB7-H3 membrane-proximal domain, predicted using AlphaFold 3, showing residues at which substitutions disrupted binding of VHH-Fcs 620, 621, 622, 633, and 636, which share an epitope in the IgV-like region, and of VHH-Fc 628, which binds the IgC-like region. Structures are shown in a reference orientation (Ref 0°) and rotated 90° about the vertical axis. Residues are shaded dark or light according to the scoring criteria described in panel (a). **(c)** Surface representation as in (b), showing residues at which substitutions disrupted binding of the benchmarking mAbs mirzotamab (purple), vobramitamab (green), and ifinatamab (blue).

Within the shared footprint of the five VHH-Fcs, Thr298 emerged as a hotspot residue, with most substitutions at this position abolishing binding (Supplementary Figure S9). The equivalent position is an alanine in the membrane-distal domain and in 2Ig, providing a molecular explanation for the preferential recognition of the membrane-proximal domain and of 4Ig by VHH-Fc 620, 621, 633 and 636. Consistent with this, VHH-Fc 622, whose footprint is more restricted and does not include Thr298, showed little domain discrimination by SPR.

Epitope mapping of clinical benchmarking mAbs revealed distinct binding profiles across the molecule. Mirzotamab and vobramitamab recognized epitopes in the upper, loop region of the IgV-like domain, on opposite faces of the molecule. The vobramitamab epitope partially overlapped with the VHH-Fc-targeted region. In contrast, ifinatamab bound the central region of the IgC-like domain and partially overlapped with the epitope recognized by VHH-Fc 628 (Figure 3c).

We extended the mapping of VHH-Fc 620, VHH-Fc 636, and ifinatamab to the membrane-distal domain. As expected, their epitopes localized to regions homologous to those identified in the membrane-proximal domain (Supplementary Figure S10, S11). Notably, the VHH-Fcs showed greater sensitivity to substitutions at residues spatially distant from the mapped epitope. This increased sensitivity may reflect their lower affinity for the membrane-distal domain and/or a greater dependence on the structural integrity of this domain.

Together, DMS resolved the VHH-Fc panel into two epitope groups corresponding to the bins defined by SPR, localizing VHH-Fc 620, 621, 622, 633 and 636 to the IgV-like region overlapping the vobramitamab epitope, and VHH-Fc 628 to the IgC-like region overlapping that of ifinatamab. The residue-level maps additionally account for the domain and isoform selectivity observed by SPR.

### VHH-Fc 620 exhibits superior B7-H3-dependent receptor internalization

Having localized the epitopes of the VHH-Fc panel, we next asked whether binders sharing a footprint also behave similarly on cells. Because the activity of the linker-drug used in this study depends on delivery of the payload to acidic endolysosomal compartments, where cathepsins cleave the linker and release the payload, we compared the internalization capacity of the VHH-Fcs using a pH-sensitive label that is non-fluorescent at extracellular pH and becomes fluorescent on trafficking to endosomes and lysosomes (pH ∼4.5–5.5) (Figure 4a). Fluorescence therefore reports internalized rather than surface-bound binders. Binders were tested at 13.5 nM and 6.75 nM on RD and RH30 WT cells, with RH30 KO cells and an isotype control included to establish hB7-H3 dependence. Ifinatamab was included as a benchmarking mAb, to which endpoint values were normalized.

**Figure 4.**
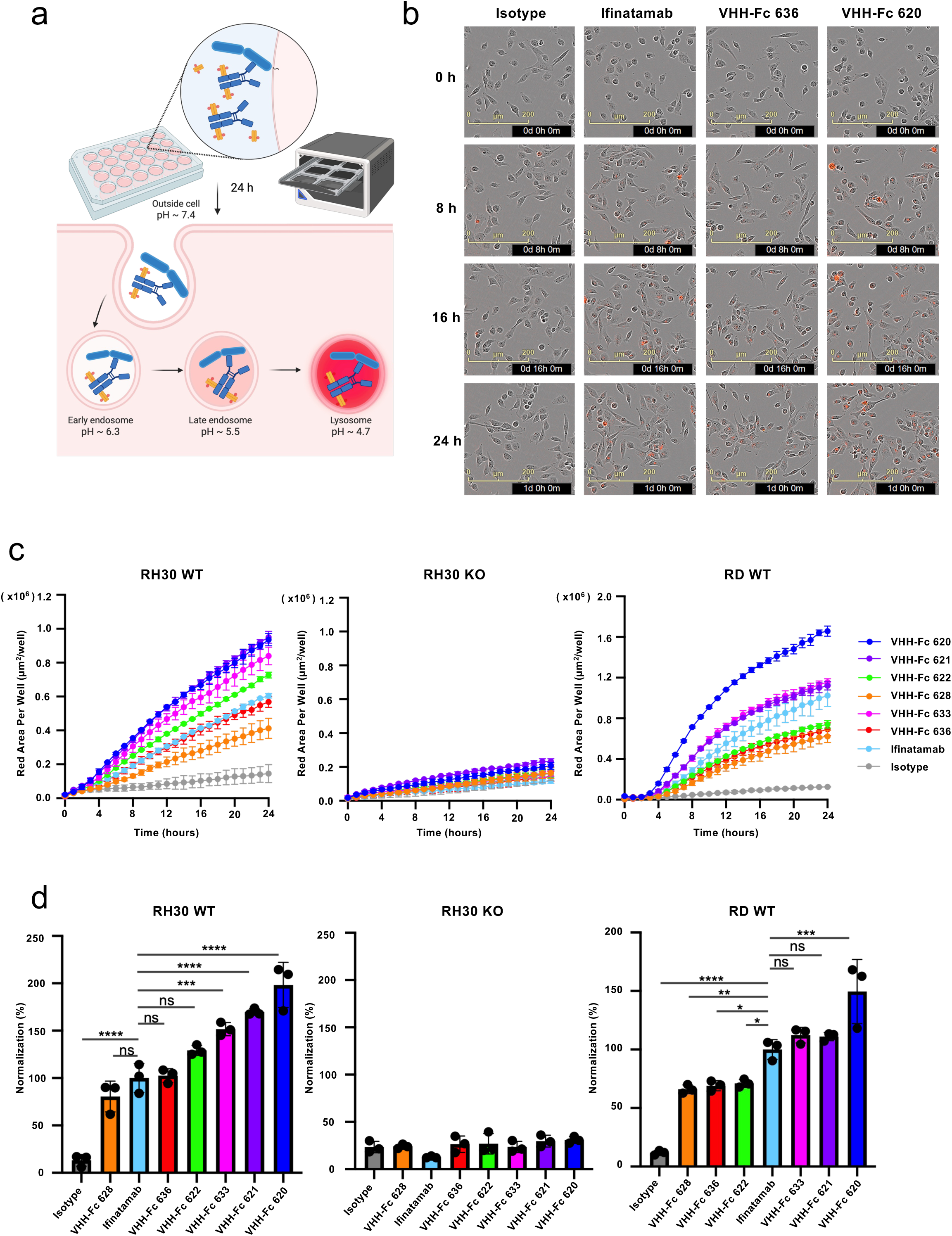
Real-time analysis of VHH-Fc internalization using the Incucyte live-cell imaging system. **(a)** Schematic of VHH-Fc internalization following binding to hB7-H3 at the cell surface. Binders were labelled with a pH-sensitive dye that is non-fluorescent at extracellular pH and becomes fluorescent on trafficking to acidic endosomal and lysosomal compartments. Created in BioRender. Sudareva, M. (2026) https://BioRender.com/rn4aw4a **(b)** Representative Incucyte images (phase contrast with red fluorescence overlay) of RH30 wild-type (WT) cells at 0, 8, 16, and 24 h after treatment with 13.5 nM isotype control, ifinatamab, VHH-Fc 636, and VHH-Fc 620. Red fluorescence indicates intracellular accumulation of labeled binders, reflecting internalization over time. **(c)** Quantitative kinetic analysis of VHH-Fc and ifinatamab internalization at 13.5 nM in RH30 knock-out (KO), RH30 WT, and RD WT cells, measured as red object area per well over 24 h using the Incucyte system. Data are shown as mean ± SD from one representative assay of three independent experiments with similar results, each including three technical replicates. **(d)** Endpoint quantification at 24 h of the data shown in (c), normalized to ifinatamab in the WT cell lines. Bars represent mean ± SD from three independent experiments, each including three technical replicates. Statistical significance was determined by one-way ANOVA with Dunnett’s multiple comparisons test against ifinatamab (* p < 0.05, ** p < 0.01, *** p < 0.001, **** p < 0.0001; ns, not significant) in GraphPad Prism 10.

Live-cell imaging over 24 h revealed progressive intracellular accumulation of all VHH-Fcs in both RD and RH30 WT cells, with fluorescence signal increasing over time, consistent with time-dependent internalization (Figure 4b-c, Supplementary Figure S12a-b). The kinetic curves also revealed a clear concentration-dependent effect, with 13.5 nM VHH-Fcs yielding faster and greater signal accumulation than 6.75 nM across the tested binders. In contrast, minimal fluorescence was detected in RH30 KO cells for all conditions, comparable to the isotype control, confirming that uptake in the WT cells was B7-H3-dependent.

Marked differences in internalization efficiency were observed among the VHH-Fcs, with VHH-Fc 620 consistently generating the highest intracellular fluorescence signal in both WT cell lines and at both concentrations, exceeding that observed for all other VHH-Fcs and ifinatamab (Figure 4c, Supplementary Figure S12b). Endpoint quantification at 13.5 nM confirmed this ranking. In RH30 WT cells, VHH-Fc 620, 621 and 633 achieved significantly greater internalization than ifinatamab, whereas VHH-Fc 636 was comparable to ifinatamab and VHH-Fc 628 showed the weakest internalization (Figure 4d). In RD WT cells, VHH-Fc 620 again showed the highest internalization of all VHH-Fcs, significantly exceeding ifinatamab, while VHH-Fc 628, VHH-Fc 636, and VHH-Fc 622 formed a lower cluster with internalization below that of ifinatamab. At 6.75 nM, the overall trend was largely preserved, with VHH-Fc 620 remaining the highest-internalizing candidate and VHH-Fc 628 the weakest in both cell lines (Supplementary Figure S12c).

In summary, the internalization studies demonstrated that epitope and affinity do not fully determine internalization, as exemplified by VHH-Fc 620 and VHH-Fc 636, which share a footprint and comparable binding profiles yet differ markedly in uptake. VHH-Fc 620 achieved the most efficient and reproducible hB7-H3-mediated internalization across both cell lines and both concentrations, consistently outperforming ifinatamab.

### ADC potency does not follow from internalization efficiency

Having characterized the VHH-Fc panel by affinity, epitope and internalization, we next asked whether these properties predict their cytotoxic activity in the ADC format. VHH-Fc 620 was selected as the lead candidate, combining domain- and isoform-selective binding with the most efficient internalization of the panel, and was benchmarked against ifinatamab conjugated to the same linker and payload.

Both VHH-Fc 620 and ifinatamab were conjugated to the topoisomerase I inhibitor exatecan through a protease-cleavable MC-GGFG-PAB linker (MedChemExpress) (Figure 5a). Exatecan was chosen as a payload of the same class as DXd, the exatecan derivative used in ifinatamab deruxtecan, which has shown clinical activity across hB7-H3-expressing solid tumors^21^. Exatecan itself has established utility as an ADC payload in preclinical studies^24,25^. The MC-GGFG-PAB linker was selected to remain stable under assay conditions while permitting efficient lysosomal release following receptor-mediated internalization. Both conjugates were prepared with the same linker and payload, allowing a direct comparison of the targeting domain. Conjugate quality was confirmed by high-performance liquid chromatography (HPLC) and mass spectrometry, giving drug-to-antibody ratios (DAR) of 1.9 for VHH-Fc 620–exatecan and 2.25 for ifinatamab–exatecan (Supplementary Figure S13a–b). Conjugation did not affect binding to 4Ig hB7-H3, with comparable affinities for VHH-Fc 620 and VHH-Fc 620–exatecan (KD = 3.51 and 3.35 nM), and for ifinatamab and ifinatamab–exatecan (KD = 4.82 and 4.85 nM) (Figure 5b, Supplementary Table S3).

**Figure 5.**
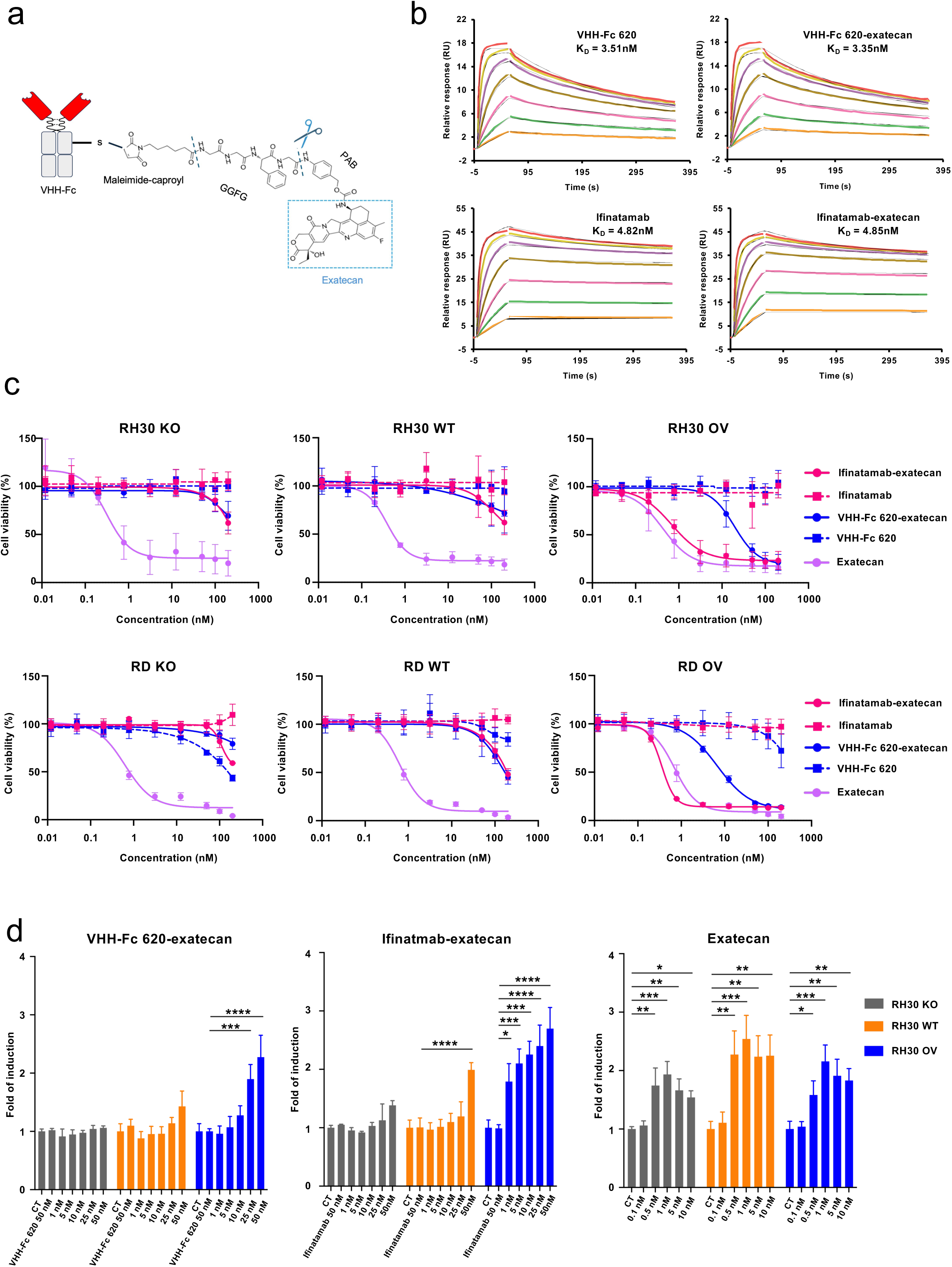
Exatecan conjugation preserves B7-H3 binding affinity and mediates B7-H3-dependent cytotoxicity in sarcoma cell lines. **(a)** Schematic representation of the VHH-Fc 620–exatecan conjugate. Exatecan was conjugated to the VHH-Fc via a cleavable maleimide-caproyl–GGFG–PAB linker. Conjugation was performed by MedChemExpress; see Supplementary Figure S13 for characterization. **(b)** Representative SPR sensograms showing binding of unconjugated VHH-Fc 620 and ifinatamab, as well as VHH-Fc 620–exatecan and ifinatamab–exatecan, to recombinant 4Ig hB7-H3. KD values are indicated on each sensorgram. Data shown are from one representative experiment of three independent experiments with similar results. **(c)** Concentration–response curves showing the effects of VHH-Fc 620–exatecan, ifinatamab–exatecan, the corresponding unconjugated binders, and free exatecan on cell viability in wild-type (WT), B7-H3-knockout (KO), and B7-H3-overexpressing (OV) RH30 and RD cells, measured after 120 h of treatment using CellTiter-Glo® 2.0. Cell viability was normalized to that of untreated controls. Data are presented as mean ± SD from three independent experiments, each performed in technical triplicate. Concentration– response curves were fitted by non-linear regression using the variable-slope four-parameter logistic model ([inhibitor] vs. response) in GraphPad Prism 10. IC₅₀ values are summarized in Supplementary Figure S15b. **(d)** Induction of apoptosis by B7-H3-targeting VHH-Fc 620–exatecan, ifinatamab–exatecan and free exatecan in RH30 cells. B7-H3 knockout (RH30 KO), wild-type (RH30 WT), and B7-H3-overexpressing (RH30 OV) cells were treated with VHH-Fc 620–exatecan, ifinatamab–exatecan, the corresponding unconjugated binders or free exatecan at the indicated concentrations for 96 h. Caspase-3/7 activity was measured using the Caspase-Glo® 3/7 assay and is presented as fold induction relative to the corresponding untreated control (CT). Data are shown as mean ± SD from 3 independent experiments. Statistical significance was determined by one-way ANOVA with Dunnett’s multiple comparisons test (* p < 0.05, ** p < 0.01, *** p < 0.001, **** p < 0.0001) in GraphPad Prism 10.

Cytotoxic activity was evaluated in four human sarcoma cell lines (RD, RH30, Saos-2 and MG63) expressing WT levels of B7-H3, together with the 4Ig hB7-H3-overexpressing (OV) and B7-H3 KO derivatives of each (Supplementary Figure S14a–b). Both ADCs showed minimal cytotoxicity in KO cells, indicating that cytotoxicity was largely B7-H3-dependent (Figure 5c, Supplementary Figure S15a–b). In contrast, free exatecan was equally cytotoxic irrespective of hB7-H3 expression (IC₅₀ 0.31–1.13 nM), consistent with its non-targeted mechanism of action. In WT cells, both ADCs had only modest effects on viability, and IC₅₀ values could not be determined within the tested concentration range (up to 200 nM). In the OV models, both ADCs were substantially more active, with IC₅₀ values of 0.35–3.36 nM for ifinatamab–exatecan and 3.03–31.27 nM for VHH-Fc 620–exatecan. Cytotoxicity correlated with the fold induction of caspase-3/7 activity in RH30 cells, which increased for both ADCs only in the OV line and for free exatecan irrespective of B7-H3 expression (Figure 5d).

In summary, the activity of both ADCs scaled with hB7-H3 expression, whereas free exatecan killed all cell lines irrespective of B7-H3 expression. This indicates that cytotoxicity was determined by hB7-H3-dependent payload delivery rather than the intrinsic sensitivity of the cell lines to exatecan. Furthermore, although both ADCs showed the same pattern of hB7-H3-dependent cytotoxicity, ifinatamab-exatecan was consistently 3-to 29-fold more potent than VHH-Fc 620-exatecan across the four OV models. This difference was not attributable to binding affinity, which was comparable between the two constructs, nor to internalization, where VHH-Fc 620 outperformed ifinatamab, and is unlikely to be explained by the modest difference in DARs. Because the OV lines express the 4Ig isoform, which is recognized equally well by VHH-Fc 620 and ifinatamab, isoform selectivity does not account for the difference either. Hence, the events following internalization will need to be further investigated to understand the differential ADC activity.

## Discussion

hB7-H3 is among the most actively pursued targets in solid tumor oncology, yet its mAbs have undergone limited comparative characterization, and the duplicated architecture of its ECD and its soluble forms have received relatively little attention despite their potential to influence drug efficacy. In particular, which of the two IgV-IgC domain pairs an antibody engages, and whether it distinguishes the 4Ig and 2Ig isoforms, remain mostly unaddressed. Here we generated a panel of six hB7-H3-targeting VHH-Fcs and characterized them by SPR, epitope binning, domain-resolved DMS, and receptor internalization, benchmarked throughout against three clinically advanced hB7-H3 mAbs. This integrated analysis identified binders that discriminate between the membrane-proximal and membrane-distal IgV-IgC domains of hB7-H3 and between the 4Ig and 2Ig isoforms. VHH-Fc 620 was selected as a lead candidate, which we conjugated to exatecan and compared directly with ifinatamab conjugated to the same linker and payload.

Despite the high sequence identity of the IgV-IgC domains within the 4Ig and 2Ig isoforms, most VHH-Fcs preferentially bound the membrane-proximal domain, and bound the membrane-distal domain and 2Ig with substantially lower affinity. The benchmarking mAbs displayed no such discrimination, binding both isolated domains and both isoforms with comparable affinity. Hence, subtle sequence and structural differences are sufficient for the VHH-Fcs to discriminate between the duplicated domains and the two isoforms.

The preferential recognition of the membrane-proximal domain prompted us to define the epitopes of the panel at residue level. We performed epitope binning to classify binders by competition, and DMS to identify the contribution of individual surface residues to binding, together providing a more complete picture of binder-antigen interaction. Binning grouped the panel into two major bins, one containing five VHH-Fcs as well as vobramitamab and a second containing VHH-Fc 628 and ifinatamab, with mirzotamab in an adjacent third bin. DMS added resolution in two respects. Although vobramitamab and the five VHH-Fcs share a bin, their footprints only partially overlap, placing the VHH-Fcs on a surface adjacent to rather than coincident with the vobramitamab epitope. DMS also independently supported the assignment of VHH-Fc 628 to the ifinatamab bin, with both epitopes mapping to the core of the membrane-proximal IgC-like domain. This is relevant because the VHH-Fc 628 binning experiment was performed on full-length 4Ig hB7-H3 rather than the isolated domain.

DMS also provided a molecular explanation for the domain preference. Thr298 emerged as a hotspot residue within the footprint of the dominant VHH-Fc epitope, with most substitutions at this position abolishing binding, and is replaced by alanine in both the membrane-distal domain and 2Ig, accounting for the reduced binding to each. Although we did not screen specifically for binders engaging this position, Thr298 is one of the few surface residues that differ between the two domain pairs and that antibodies could engage, and therefore one of the few residues through which isoform selectivity can be achieved. VHH-Fc 622, whose footprint is more restricted and does not include Thr298, showed correspondingly little domain discrimination.

Whether isoform selectivity is desirable depends on the therapeutic context. Selective binders could reduce sequestration by circulating soluble hB7-H3, which is present in healthy individuals and elevated in patients with osteosarcoma and other malignancies at concentrations comparable to the affinities measured here^6,7,26^. The alternatively spliced soluble form corresponds to the 2Ig ECD, with four extra residues at the C-terminus^6^, and would be poorly recognized by most of the VHH-Fcs, although a shed 4Ig ECD would retain the membrane-proximal domain and be bound. This rationale has recently been pursued with MIL33B, an antibody with 8-fold selectivity for human 4Ig over 2Ig developed for radioligand therapy^8^. The concern is greatest for therapeutic modalities administered at low protein mass, where circulating B7-H3 can consume a substantial fraction of the dose, including radiotracers and radioimmunotherapy, T-cell engagers, and ADCs. Conversely, binders that engage both isoforms offer broader coverage where the dominant isoform is unknown or variable, which may be preferable for applications less constrained by dose. The panel described here spans both behaviors, providing a set of tools with which these questions can be addressed.

As shown for HER2-targeting antibodies, epitope localization can influence receptor internalization and intracellular trafficking^13^. Affinity contributes as well, with variants against a single HER2 epitope showing that it affects both cellular uptake and ADC potency in a manner dependent on antigen density and payload^27^. Internalization itself is therefore commonly used as a selection criterion for ADC leads^27,28^. All VHH-Fcs underwent hB7-H3-dependent internalization, indicating that preferential targeting of the membrane-proximal domain is compatible with receptor-mediated uptake. Within the main epitope bin, however, uptake varied several-fold. VHH-Fc 620 accumulated most in both cell lines, while VHH-Fc 636, despite near-identical epitope and affinity, internalized substantially less. VHH-Fc 622 and VHH-Fc 628 showed the lowest uptake overall, which may reflect the faster dissociation of VHH-Fc 622 and the weaker affinity of VHH-Fc 628, but neither explanation extends to VHH-Fc 636. VHH-Fc 622 and VHH-Fc 636 also ranked differently between RH30 and RD cells, suggesting that cell-intrinsic factors contribute alongside the properties of the binder itself. Internalization is therefore a distinct property rather than a consequence of binding affinity and epitope, and led us to select VHH-Fc 620 for conjugation.

Both ADCs, VHH-Fc 620-exatecan and ifinatamab-exatecan, showed B7-H3-dependent cytotoxicity, with IC_50_ values determinable only in the OV lines and minimal activity in the WT and KO cells at concentrations up to 200 nM, whereas free exatecan was equally cytotoxic irrespective of antigen expression. This is consistent with antigen density as an important determinant of ADC efficacy^29,30^, and confirms that killing was antigen-directed. However, the superior internalization of VHH-Fc 620 did not translate into greater cytotoxicity. Ifinatamab-exatecan was 3-to 29-fold more potent than VHH-Fc 620-exatecan across all four OV lines, despite comparable binding affinity and lower cellular uptake. Internalization is therefore necessary but not sufficient for potent ADC activity, which also depends on intracellular trafficking, lysosomal processing and payload release, each of which determines how much free drug reaches its target^31,32^.

Two technical differences may contribute. The conjugates differed modestly in DAR (1.9 versus 2.25) and in purity (89.6 % versus 98.2 %), although neither accounts for a difference of this magnitude on its own. Whether the VHH-Fc format is routed differently after internalization remains an open question. As neither ADC was active at endogenous levels of hB7-H3, we did not proceed to *in vivo* efficacy studies at this stage. Because both conjugates showed this limitation, it is unlikely to reside in the binder itself. The low DARs are one contributing factor, and for the VHH-Fcs this partly reflects a structural constraint, since the two hinge disulfides provide only four conjugation sites compared with eight in an IgG1. Site-specific conjugation strategies that do not rely on interchain cysteines would therefore be required to raise payload loading. Resolving this is worthwhile, as the smaller VHH-Fc format may offer faster tumor accumulation and improved penetration relative to full-length IgG^33,34^.

In conclusion, we generated a panel of hB7-H3-targeting VHH-Fcs and characterized them by binding kinetics, domain-resolved epitope mapping, receptor internalization and functional ADC evaluation. Most of the panel discriminated between the two IgV-IgC domain pairs of 4Ig hB7-H3 and between the 4Ig and 2Ig isoforms. At residue level, the selectivity maps to Thr298, which differs between the two domains and 2Ig. The clinical benchmarking mAbs did not display this selectivity, although it may be advantageous where circulating soluble B7-H3 competes for binding. Furthermore, the functional characterization showed that neither affinity nor epitope predicted internalization, and that internalization in turn did not predict ADC potency. VHH-Fc 620 was selected as the lead candidate on this basis and, although further optimization of the conjugate and validation *in vivo* are needed, provides a starting point for isoform-selective B7-H3 targeting.

## Materials and methods

### Generation and selection of hB7-H3 specific VHH

VHHs were generated by llama immunization and selected by phage display as previously described^35^. Briefly, immunization and VHH phage library construction were performed by QVQ B.V. (Utrecht, the Netherlands) with in-house expressed and purified antigens (see below). Two independent immunization campaigns, each involving two llamas and distinct immunization strategies (Figure 1a), were conducted. In the first campaign, two llamas received four subcutaneous injections of a mixture of recombinant 4Ig hB7-H3 and 2Ig mB7-H3 ECDs to generate cross-reactive VHHs. In the second campaign, two llamas received six subcutaneous injections of recombinant 4Ig hB7-H3 ECD to generate hB7-H3-specific VHHs with enhanced binding affinity. Following immunization, PBMCs were collected, and total RNA was isolated from lymphocytes. VHH-encoding sequences were then amplified to construct immune phage display libraries. VHHs were displayed on M13 bacteriophage, and two rounds of biopanning were performed using human 4Ig hB7-H3 ECD. Phage ELISA was used to identify VHHs specific for 4Ig hB7-H3. Selected candidates were subsequently expressed as described below for further characterization.

### Production and purification of VHH-Fcs and hB7-H3 ECDs

All constructs were expressed by transient transfection in Expi293F cells (Thermo Fisher Scientific) according to the manufacturer’s protocol.

To produce the VHH-Fcs for functional studies, the coding sequences of VHHs 620, 621, 622, 628, 633, and 636 were fused to the human IgG1 Fc region and cloned into the pcDNA3.1 expression vector (Thermo Fisher Scientific). Each VHH contained an N-terminal HA signal peptide (MKTIIALSYIFCLVFA), and a C-terminal 3C protease cleavage site (LEVLFQ/GP), the human IgG1 Fc-tag, and a 6×His-tag. Following expression in Expi293F cells, the VHH–Fc proteins were purified using CaptivA Protein A Affinity Resin (Repligen) and further purified by SEC on a Superdex 200 column (Cytiva) equilibrated with running buffer containing 20 mM HEPES pH 7.3 and 500 mM NaCl.

To produce recombinant antigens for llama immunization, the 4Ig hB7-H3 ECD (residues 29-457) was cloned into the pcDNA3.1 expression vector (Thermo Fisher Scientific) with an N-terminal kappa light chain signal peptide (MDMRVPAQLLGLLLLWLRGAR), and a C-terminal 3C protease cleavage site (LEVLFQ/GP), the human IgG1 Fc-tag, and a 6×His-tag. Following expression in Expi293F cells, B7-H3 ECD-Fc protein was purified using CaptivA Protein A Affinity Resin (Repligen). Following elution with Pierce™ IgG elution buffer, the C-terminal tag was cleaved with PreScission Protease (Cytiva) and cleared with Ni Sepharose Excel resin (Cytiva). The B7-H3 ECD was further purified by SEC on a Superdex 200 column (Cytiva) equilibrated with running buffer containing 20 mM HEPES pH 7.3 and 500 mM NaCl. The 2Ig mB7-H3 ECD (residues 29-238) was cloned into the pcDNA3.1 expression vector (Thermo Fisher Scientific) with an N-terminal kappa light chain signal peptide (MDMRVPAQLLGLLLLWLRGAR), and a C-terminal biotin acceptor peptide (BAP)-tag (GLNDIFEAQKIEWHE) and a 6xHis-tag. Following expression in Expi293F cells (Thermo Fisher Scientific), the mB7-H3 ECD was purified using Ni Sepharose excel (Cytiva), and SEC on a Superdex 200 column (Cytiva) equilibrated with running buffer containing 20 mM HEPES pH 7.3 and 500 mM NaCl.

To produce hB7-H3-Fc for ELISA, the full-length 4Ig hB7-H3 (residues 29-457) was cloned into the pcDNA3.1 expression vector (Thermo Fisher Scientific) with an N-terminal HA signal peptide (MKTIIALSYIFCLVFA), and a C-terminal 3C protease cleavage site (LEVLFQ/GP), the human IgG1 Fc-tag, and a 6×His-tag. Following expression in Expi293F cells (Thermo Fisher Scientific), the hB7-H3–Fc proteins were purified using CaptivA Protein A Affinity Resin (Repligen) and further purified by SEC on a Superdex 200 column (Cytiva) equilibrated with running buffer containing 20 mM HEPES pH 7.3 and 500 mM NaCl.

To produce recombinant B7-H3 ECD variants for functional assays, the full-length 4Ig hB7-H3 (residues 29-457), the membrane-distal domain of 4Ig (residues 29–238), the membrane-proximal domain of 4Ig (residues 243–456), and the 2Ig (residues 29–238), containing a C-terminal 3C protease cleavage site (LEVLFQ/GP), a biotin acceptor peptide (BAP)-tag (GLNDIFEAQKIEWHE) and a 6xHis-tag were cloned into the pcDNA3.1 expression vector (Thermo Fisher Scientific). Following expression in Expi293F cells (Thermo Fisher Scientific), the B7-H3 ECDs were purified using Ni Sepharose excel (Cytiva) affinity purification, and Superdex 75 and 200 (Cytiva) SEC column equilibrated with running buffer containing 20 mM HEPES pH 7.3 and 150 mM NaCl. Enzymatic biotinylation was performed in 50 mM bicine pH 8.3, 10 mM ATP, 10 mM Mg(OAc)_2_, 0.5 mM d-biotin with recombinant GST-BirA ligase overnight at 4 °C, and proteins were subsequently re-purified by SEC on a Superdex 75 or 200 column (Cytiva) to remove excess biotin.

Infinitamab was purchased from MedChemExpress (HY-P99367, lot#257463), mirzotamab from MedChemExpress (HY-P99740, lot#271609), and vobramitamab from MedChemExpress (HY-P99101, lot#494240).

### Thermal stability assay

Protein thermal stability was measured by differential scanning fluorimetry using the Protein Thermal Shift™ Dye Kit (Thermo Fisher Scientific, Cat. No. 4461146). The dye was diluted from 1000X to 8X and mixed with protein, buffer, and water to a final volume of 20 µL per well; each condition was run in four technical replicates. Samples were heated from 25 °C to 99 °C on a QuantStudio™ 12K Flex (Thermo Fisher Scientific), and Tm values were determined from the resulting melt curves using the Protein Thermal Shift™ Software (Cat. No. 4477802), which fits a Boltzmann model to the fluorescence data.

### ELISA of VHH-Fc panel

Nunc Maxisorp 96 wells plates were coated overnight at 4 °C with 100 µL per well of PBS containing 4Ig hB7-H3-Fc at a concentration of 2 µg/mL. Plates were washed three times with PBS containing 0.1 % Tween-20 (PBST) and subsequently blocked with 2 % (w/v) skim milk in PBS for 1 h at 20 °C. After three additional washes with PBST, VHH-Fcs were added at the indicated concentrations (25, 6.25, 1.56, 0.39, 0.09, 0.024, and 0.006 nM) and incubated for 1 h at 20 °C with shaking. Plates were then washed three times with PBST and incubated for 1 h at 20 °C with shaking with a rabbit anti-VHH monoclonal antibody (QVQ, QE19) diluted 1:5,000 in PBST. Following three washes with PBST, a goat anti-rabbit secondary antibody (Abcam, ab205718) diluted 1:10,000 in PBST was added and incubated for 1 h at 20 °C with shaking. After a final washing step, 100 µL of TMB substrate solution (Thermo Fisher Scientific) was added to each well. The reaction was stopped after 10 min by adding 50 µL of 1 M HCl, and absorbance was measured at 450 nm using a microplate reader (Spectrostar nano, BMG labtech).

### Cell culture

Human osteosarcoma and rhabdomyosarcoma cell lines were maintained in an incubator at 37 °C, 95 % humidity and 5 % CO₂. RD cells were cultured in Dulbecco’s Modified Eagle Medium (DMEM) (Gibco) supplemented with 10 % fetal bovine serum (FBS) (Sigma-Aldrich), 100 U/mL penicillin (Gibco) and 100 μg/mL streptomycin (Gibco). Saos-2 cells were maintained in DMEM (Gibco) supplemented with 20 % FBS (Sigma-Aldrich), 100 U/mL penicillin (Gibco) and 100 μg/mL streptomycin (Gibco). MG63 and RH30 cells were cultured in Roswell Park Memorial Institute (RPMI) 1640 medium (Gibco) supplemented with 10 % FBS (Sigma-Aldrich), 100 U/mL penicillin (Gibco) and 100 μg/mL streptomycin (Gibco). Cells were passaged upon reaching approximately 70-90 % confluency using standard cell culture procedures. Cells tested negative for mycoplasma.

### Generation of hB7-H3 knockout (KO) cell lines

hB7-H3 KO cell lines (Saos-2, RH30, and RD) were generated using CRISPR/Cas9-mediated genome editing. A guide RNA targeting human B7-H3 (5’-GUGCAGCCCUGGGAGCACUG-3’) was cloned into a lentiviral transfer vector encoding Cas9 and a puromycin resistance cassette as previously described^36^. Lentiviral particles were produced by co-transfecting HEK293T cells at 80-90 % confluency with the transfer plasmid alongside packaging plasmids (pHDM-HGPM2, pRC-CMV-rev1b, pHDM-tat1b, and pHDM-G) at a ratio optimized for each transfer plasmid, using 25 kDa polyethylenimine (PEI) (Tebubio, 23966-1) as the transfection reagent. Transfection medium was replaced 18 h post-transfection with fresh DMEM (Gibco) supplemented with 10 % heat-inactivated FBS (Sigma-Aldrich), and viral supernatant was harvested at 72 h post-transfection. The supernatant was clarified by centrifugation at 500 × *g* for 5 min, filtered through a 0.45 μm membrane, aliquoted, snap-frozen in liquid nitrogen, and stored at -80 °C until use.

WT cell lines were seeded in 6-well plates and transduced with lentiviral supernatant at 1:10 and 1:50 dilution in the respective complete growth medium containing 8 μg/mL polybrene (Sigma-Aldrich) using a reverse transduction approach. After 72 h, transduced cells were selected with 2 µg/ml puromycin (Invitrogen, ant-pr) for up to 14 days, with media changes every 2-3 days, until all cells in the untransduced control well were eliminated. Antibiotic-resistant polyclonal populations were subsequently subjected to limiting dilution to isolate single-cell clones. Individual clones were screened for loss of hB7-H3 surface expression by flow cytometry using an anti-hB7-H3 antibody (clone 7-517, BD) and goat anti-mouse IgG (H+L) cross-adsorbed secondary antibody, Alexa Fluor 647 (Invitrogen, A-21235). Clones showing complete absence of B7-H3 staining were advanced for genomic confirmation. Genomic DNA was extracted and the region flanking the gRNA target site was amplified by PCR using primers (Forward 5’-GTCCAGGAGAGATGAGGCAA -3’ and Reverse 5’-ACAGGGAGGAAGCAAGCAAA -3’) designed to distinguish WT and edited alleles based on amplicon size. PCR products were resolved by agarose gel electrophoresis, and individual bands were gel-purified and subjected to Sanger sequencing. Sequencing chromatograms were aligned to the WT hB7-H3 reference sequence to characterize the indels on each allele and confirm biallelic disruption of the hB7-H3 locus in all three cell lines (Supplementary Figure S2a-c).

### FACS of VHH-Fc panel

Cells were harvested and washed twice with PBS containing 1 % BSA (PBS-B). A total of 1 × 10^5^ cells per condition were seeded into each well of a V-bottom 96-well plate and incubated at 4 °C for 30 min with either 40 nM anti–human B7-H3 antibody (clone 7-517, BD), 40 nM isotype control (human IgG1 κ, clone R1, Sino Biological), or 40 nM of VHH-Fc diluted in cold PBS-B. Following centrifugation at 350 × g for 5 min at 4 °C, supernatants were discarded and cells were washed twice with 130 µL of PBS-B. Cells were then resuspended and incubated for 30 min at 4 °C with secondary antibodies diluted 1:200 in PBS-B. Donkey anti–human IgG (H+L), cross-adsorbed DyLight 650–conjugated antibody (SA5-10129, Thermo Fisher Scientific) was used for detection of human isotype control and VHH-Fcs, while goat anti-mouse IgG (H+L), cross-adsorbed Alexa Fluor 647–conjugated antibody (A-21235, Invitrogen) was used for clone 7-517 detection. After two additional washes with 130 µL of PBS-B, cells were resuspended in 100 µL PBS-B supplemented with DAPI (1:400; ThermoFischer). Data were acquired on a CytoFLEX flow cytometer (Beckman Coulter) and analyzed using FlowJo software (FlowJo 10.10.0).

### Surface plasmon resonance (SPR) of VHH-Fc panel

The binding kinetics of VHH-Fcs to hB7-H3 were determined by SPR using a Biacore 1K system (Cytiva, Sweden) at an analysis temperature of 25 °C using HBS-EP+ running buffer (10 mM HEPES, 150 mM NaCl, 3 mM EDTA, 0.05 % v/v surfactant P20, pH 7.4). AviTag-biotinylated hB7-H3 proteins, including the 4Ig full-length, 4Ig membrane-proximal domain, 4Ig membrane-distal domain, and 2Ig constructs, were immobilized on a Series S Sensor Chip SA according to the manufacturer’s instructions. Seven 1:2 serial dilutions of the selected VHH-Fcs and benchmarking mAbs (ranging from 1.56 nM to 100 nM) were injected at a flow rate of 30 µL/min with an association phase of 60 s, followed by a 300 s dissociation phase in running buffer. After each association–dissociation cycle, the sensor surface was regenerated by injecting 2M MgCl_2_ at 30 µL/min for 300 s. SPR data were acquired using Biacore 1K Insight Control Software (Cytiva, version 5.08.18.22405) and analyzed with Biacore 1K Insight Evaluation Software (Cytiva, version 5.08.18.22102). Kinetic parameters were determined by fitting the sensorgrams to a bivalent analyte binding model which describes the interaction as a two-step process: an initial 1:1 binding event characterized by the association rate constant (k_a1_, M⁻¹·s⁻¹) and dissociation rate constant (k_d1_, s⁻¹), followed by a second, avidity-driven binding event characterized by k_a2_ (RU⁻¹·s⁻¹) and k_d2_ (s⁻¹). The apparent equilibrium dissociation constant (K_D_, M) was calculated as k_d1_/k_a1_ to reflect the functional (avidity-corrected) affinity.

### Epitope binning of VHH-Fc and mAb panel

Epitope binning of VHH-Fcs and benchmarking mAbs against hB7-H3 was performed by SPR using a Biacore 1K system (Cytiva) at an analysis temperature of 25 °C in HBS-EP+ buffer (10 mM HEPES, 150 mM NaCl, 3 mM EDTA, 0.05 % v/v surfactant P20, pH 7.4). Each VHH-Fc and benchmarking mAb was individually immobilized as capture binders on a CM5 sensor chip by amine coupling according to the manufacturer’s instructions (target immobilization level, 6,000-8,000 RU). After covalent immobilization, the membrane-proximal domain of 4Ig hB7-H3 or the full-length 4Ig hB7-H3 was injected at a flow rate of 10 µL/min for 120 s followed by injection of the second binder at a flow rate of 10 µL/min for 120 s. After each cycle, the CM5 chip was regenerated by injecting 2 M MgCl₂ at 30 µL/min for 300 s. Binning data were acquired using Insight Control Software (Cytiva, version 5.08.18.22405) and analyzed with Insight Evaluation Software using the binning extension (Cytiva, version 5.08.18.22102).

### Epitope mapping experiments

Epitopes were mapped by DMS using yeast display libraries of the membrane-proximal and membrane-distal domains of 4Ig hB7-H3. Briefly, each of the two domains of hB7-H3 (CD276; UniProt accession Q5ZPR3) was displayed independently on the surface of yeast using the corresponding coding sequence. The membrane-distal domain (amino acid residues 29–238) and the membrane-proximal domain (residues 243–456) were cloned into pCT302, positioned upstream of an HA tag (YPYDVPDYA) and Aga2P coding sequence.

Single amino acid substitution libraries were generated separately for the two domains of 4Ig hB7-H3 by splicing by overlap extension PCR (SOE-PCR) using NNK-degenerate primers (Supplementary Data 1). Following mutagenesis, the respective domain sequences were assembled and amplified by SOE-PCR. To facilitate Illumina sequencing, each domain library was subdivided into four contiguous sublibraries. For the membrane-distal domain, the sublibraries (L1 to L4) covered residues 29–92, 93–156, 157–220, and 221–238. For the membrane-proximal domain, the sublibraries (L5 to L8) covered residues 243–306, 307–370, 371–434, and 435–456.

Preparation of competent *Saccharomyces cerevisiae* EBY100 cells (ATCC MYA-4941) and yeast transformation were performed as previously described^37^. Briefly, each library was generated by gap repair cloning following electroporation of yeast cells with 1 μg of linearized vector and library DNA at a 12:1 molar ratio (insert:vector). Following transformation, cells were recovered in 20 mL SD-CAA medium. Once the culture reached an OD600 of approximately 0.3, cells were passaged and grown at 30 °C to an OD600 of 0.5–1.0 before induction by resuspension in 20 mL SG-CAA medium.

For FACS, 1 x 10^7^ induced yeast cells from each library were washed once with 1 mL of PBS containing 1 % BSA (PBS-B). Cells were then resuspended in PBS-B containing the test VHH-Fc at a concentration corresponding to its equilibrium dissociation constant (K_D_, Supplementary Figure S7, see VHH-Fc binding affinity determination by on-yeast binding assay below), using a volume sufficient to prevent ligand depletion, and incubated for 1 h at 20 °C with agitation (800 rpm). After one wash with 1 mL of PBS-B, cells were resuspended in PBS-B, containing 50 nM of a compatible anti-human B7-H3 monoclonal antibody (Supplementary Table S2), used as a conformational control, and incubated for an additional 1 h at 20 °C with agitation (800 rpm). Cells were subsequently washed twice with 1 mL of ice-cold PBS-B and stained with a mixture of secondary antibodies diluted in ice-cold PBS-B: donkey anti-human IgG (H+L) DyLight 650 (Invitrogen, SA5-10129; 1:50 dilution) to detect the VHH-Fc, and goat anti-mouse IgG (H+L)-R-PE (Invitrogen, P-852; 1:200 dilution) to detect the commercial anti-human B7-H3 monoclonal antibody. Following a 15-min incubation on ice, cells were washed twice with 1 mL of ice-cold PBS-B before FACS sorting. Yeast cells were sorted using a BD FACSAria Fusion cell sorter operated with BD FACSDiva software. Sorting gates were defined to isolate cells exhibiting abolished VHH-Fc binding while retaining binding to the conformational control antibody (Supplementary Figure S8). For each sorted population, cells corresponding to at least 100-fold the theoretical library diversity were collected and cultured separately in SD-CAA medium at 30 °C for 2 days.

Plasmid DNA was extracted from yeast populations using the Zymoprep Yeast Plasmid Miniprep II Kit (Zymo Research). Target regions were amplified by PCR, followed by a second PCR to incorporate sequencing adapters and sample-specific barcodes. NGS was performed on an Illumina iSeq 100 platform (2 × 150 bp, 300-cycle kit), generating a minimum of 200,000 reads per population. Sequencing data were analyzed using RStudio as previously described^23^. Briefly, paired-end reads were merged, and sequences with unexpected lengths, more than one mutation relative to the WT antigen sequence, or a Phred quality score <30 were excluded. Filtered DNA sequences were translated into protein sequences, which were subsequently clustered and counted to determine the enrichment ratio of each single amino acid substitution in the sorted populations relative to the unsorted library (Supplementary Data 2). Enrichment data were visualized as heatmaps representing the relative enrichment of individual mutations. A residue was classified as critical for binding when ≥50 % of the tested substitutions (≥10 of 19 possible amino acid substitutions, red dot on the heatmap) disrupted binding and as important when ≥25 % of substitutions (≥5 of 19, light-red dot on the heatmap) disrupted binding (Supplementary Figure S9).

### VHH-Fc affinity determination by on-yeast binding assays

On-yeast binding assays were performed as previously described^38^. Briefly, 1 x 10^6^ induced yeast cells expressing either the hB7-H3 membrane-distal IgV-IgC domain or the membrane-proximal IgV-IgC domain were incubated with anti-hB7-H3 VHH-Fcs or the benchmarking mAbs at concentrations ranging from 500 nM to 6 pM in a volume sufficient to prevent ligand depletion. Samples were incubated for 2 h at 20 °C with agitation (800 rpm). Cells were then washed twice with 1 mL of ice-cold PBS containing 1 % BSA (PBS-B) and resuspended in ice-cold PBS-B containing donkey anti-human IgG (H+L) DyLight 650 (Invitrogen, SA5-10129; 1:50 dilution). After a 15-min incubation on ice, cells were washed twice with 1 mL of ice-cold PBS-B and analyzed using an Accuri C6 flow cytometer. Flow cytometry data were processed with FlowJo v10.10.0, and equilibrium dissociation constants were determined by nonlinear regression analysis using GraphPad Prism 10.

### VHH-Fc and mAb internalization assay

Live-cell imaging experiments were performed with an Incucyte S3 system (Essen Biosciences). RH30 WT, RH30 KO and RD WT cells were seeded at 7,500 cells/well in 50 μL of culture medium in 96-well plates (Corning) and incubated overnight before treatment (37 °C, 95 % humidity, 5 % CO₂). The following day, VHH-Fcs, ifinatamab and an isotype control (human IgG1κ, clone R1, Sino Biological) were diluted in cell culture medium and incubated with 5 μg/mL Incucyte human FabFluor-pH Red antibody labeling reagent (Sartorius, Cat. 4722) for 20 min at room temperature. A fixed dye concentration was used rather than a fixed dye-to-antibody mass ratio, so that all binders were labelled at equivalent molar stoichiometry despite the difference in molecular weight between the VHH-Fc constructs and full-length IgG.

Labelled binders were added to the cells at final concentrations of 13.5 nM and 6.75 nM in a final well volume of 100 μL, and plates were immediately transferred to the Incucyte S3 live-cell analysis system and imaged for phase contrast and red fluorescence with a 20× objective every hour for 24 h. Each condition was run in triplicate wells, with two micrographs captured and analyzed per well per time point, and the assay was performed in three independent experiments.

Images were analyzed using Incucyte S3 software version 2025B. Red fluorescence was detected by surface fit segmentation with a mean intensity filter between 0.1 and 4 red calibrated units (RCU), a threshold of 0.2 RCU, and a minimum area filter of 750 μm². Total red fluorescence area (μm²/well) was plotted against incubation time after background subtraction. For endpoint quantification, the total red area at 24 h was normalized to the value obtained for ifinatamab in the corresponding wild-type cell line and expressed as a percentage. Statistical significance was assessed by one-way ANOVA with Dunnett’s multiple comparisons test against ifinatamab using GraphPad Prism 10.

### Antibody–drug conjugates

VHH-Fc 620 and ifinatamab were conjugated to exatecan through a protease-cleavable MC-GGFG-PAB linker by MedChemExpress. Conjugation, purification, and characterization by HPLC and intact mass spectrometry were performed by the supplier; representative data are shown in Supplementary Figure S13.

### Generation of B7-H3 overexpressing cell lines

RD, RH30, MG63 and Saos-2 KO cell lines were transduced with a lentiviral vector encoding hB7-H3 (provided by C.R.). Cells were cultured in complete growth medium seeded at a density of 6 × 10^5^ cells per well until they reached 80-90 % confluence in 6-well plates 24 h prior to transduction. The next day, cells were washed once with PBS and incubated with lentiviral supernatant containing polybrene (8 μg/mL). Plates were sealed with parafilm and centrifuged at 2400 rpm and 32 °C for 30 min, followed by incubation at 37 °C for 30 min. Viral supernatant was then replaced with fresh complete growth medium. Following expansion, transduced RD, MG63 and Saos-2 were single-cell sorted, and RH30 cells were bulk-sorted on a Sony SH800S cell sorter to generate hB7-H3 overexpressing cell lines. The extent and uniformity of hB7-H3 overexpression in each cell line were assessed by flow cytometry.

### Cell viability assay

Cell viability was assessed using the CellTiter-Glo® 2.0 Luminescent Cell Viability Assay (Promega, G9242). RD, MG63, RH30, and Saos-2 cells were seeded in flat-bottom 96-well plates at 1,000, 1,000, 2,000, 2,000, cells per well, respectively in 100 μL of complete growth medium and allowed to adhere overnight in the incubator at 37 °C, 5 % CO_2_, and 95 % humidity. The following day, medium was removed and the cells were treated with increasing concentration of VHH-Fc 620-exatecan, ifinatamab-exatecan, VHH-Fc 620, ifinatamab and exatecan for 120 h. After treatment, plates were acclimated to room temperature for 30 min then treated with the equal volume of CellTiter-Glo® 2.0. Plates were shaken for 20 min at room temperature. Luminescence was measured using a CLARIOstar microplate reader (BMG LABTECH), and values were corrected against medium-only background controls. IC_50_ values were determined by nonlinear regression analysis using GraphPad Prism 10.

### Cell apoptosis assay

Apoptosis was assessed in RH30 wild-type (WT), knockout (KO), and overexpression (OV) cells using the Caspase-Glo® 3/7 Assay (Promega, G8091). Cells were seeded at 4,000 cells per well in 100 µL of culture medium in 96-well plates and incubated overnight at 37 °C with 5 % CO₂. The medium was then replaced with 100 µL of fresh medium containing serial concentrations of VHH-Fc 620–exatecan, ifinatamab– exatecan, VHH-Fc 620, ifinatamab, or free exatecan. Untreated cells served as controls. After 96 h, plates were equilibrated to room temperature for 30 min. Next, 50 µL of medium was removed and replaced with 50 µL of Caspase-Glo® 3/7 Reagent. Plates were shaken at 400 rpm for 30 s and incubated at room temperature for 1 h. Subsequently, 60 µL from each well was transferred to a white 96-well plate, and luminescence was measured using a CLARIOstar microplate reader. Background-corrected luminescence was expressed as fold induction relative to untreated controls. Three independent biological experiments were performed, each with three technical replicates. Data are presented as mean ± SD and were analyzed using one-way ANOVA with Dunnett’s multiple comparisons test in GraphPad Prism 10.

## Author contributions

F.L.H.v.R., and K.v.N. produced protein for llama immunization. F.L.H.v.R., K.v.N., W.T., M.F., B.C., and C.Y.J. performed the phage display selections. X.F., B.C., S.S.G., W.T., M.F., M.C., and B.Z. produced proteins for functional characterization. B.C. and X.F. performed the biophysical characterization and deep mutational scanning, including sequencing and computational analysis. X.F., B.C., S.S.G., A.S.B., M.S., and M.F. performed the cell-based characterization, including internalization and ADC cytotoxicity assays. S.S.G., R.Mo., R.Me., and A.K. generated the B7-H3 knockout and overexpressing cell lines. B.C., X.F., S.S.G., and C.Y.J. prepared the figures and wrote the manuscript. J.H.M.M., C.R., and R.Me. supervised specific parts of the work. C.Y.J. conceived the study, secured funding, and supervised the work. All authors reviewed, edited, and approved the final version.

## Acknowledgments

The presented work was supported by the Foundation Children Cancer Free (KiKa grant 392 and 462 to C.Y.J.), Oncode Accelerator, a Dutch National Growth Fund project under grant number NGFOP2201 (to C.Y.J.) and KiKa core funding (to C.Y.J.). W.T. acknowledges financial support from the Study Abroad Program by the Government of Shandong Province (grant no. 201902012). We acknowledge the Utrecht Sequencing Facility (USEQ) for providing sequencing service and data. USEQ is subsidized by the University Medical Center Utrecht and The Netherlands X-omics Initiative (NWO project 184.034.019). We thank Dr. Christin Radon (Cytiva) and Dr. Leon Kraakman (Cytiva) for helpful discussions about the SPR affinity and epitope binning experiments, and the Flow Cytometry and Cell Sorting Facility of the Faculty of Veterinary Medicine at Utrecht University and the Princess Máxima Center for support.

## Data availability

Sequencing reads from the deep mutational scanning experiments were processed as previously described^23^, and the resulting enrichment ratios, which underlie all heatmaps shown, are provided in Supplementary Data 2. All other data supporting the findings of this study are available from the corresponding author upon reasonable request. The amino acid sequences of the VHHs described here are subject to a pending patent application and are available from the corresponding author under a material transfer agreement.

## Declaration of the use of Gen AI

In the preparation of this manuscript, the authors used Claude (Anthropic; Claude Opus 5 and Claude Opus 5.5) to assist with language editing and structuring of the text. All suggestions were reviewed and verified by the authors, who edited the text as needed and take full responsibility for the content of the publication.

## Supplementary Information

**Supplementary Table S1**

Binding affinities (KD) for the VHH-Fc and mAbs against 4Ig hB7-H3, the membrane-proximal and membrane-distal IgV-IgC domains of hB7-H3, and 2Ig hB7-H3, as determined by SPR. Kinetic parameters were determined by fitting the sensorgrams to a bivalent analyte binding model. Data are mean ± SD of three independent experiments.

**Supplementary Table S2**

Conformational control mAbs used for epitope mapping cell sorting of the VHH-Fc panel (a) and the mAb panel (b): 7-517 (BD Pharmingen™ Purified Mouse Anti-Human CD276, Cat. No. 565828) and MIH-42 (BioLegend Purified anti-human CD276, Cat. No. 50-168-631).

**Supplementary Table S3**

Binding affinity (K_D_) of VHH-Fc620, VHH-Fc620_(MC-GGFG-PAB)*_*Exatecan, Ifinatamab and Ifinatamab _(MC-GGFG-PAB)_Exatecan against 4Ig hB7H3 determined by SPR. Kinetic parameters were determined by fitting the sensorgrams to a bivalent analyte binding model. Data are mean ± SD of three independent experiments.

