## Supplementary Figures for "Domain-resolved epitope mapping of B7-H3-targeting VHH-Fcs defines isoform-selective binders for antibody–drug conjugate development"

Supplementary Figure S1

a

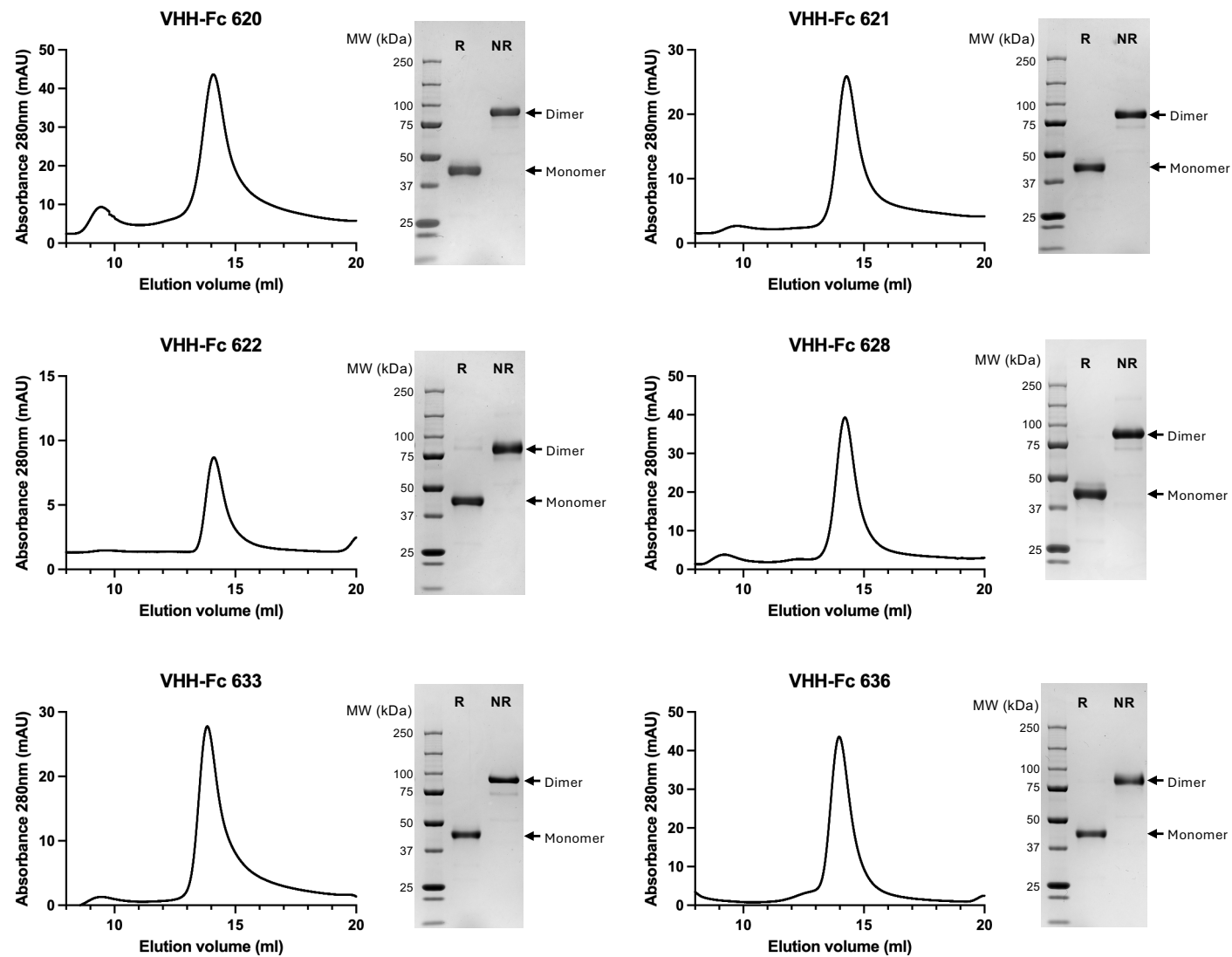

b

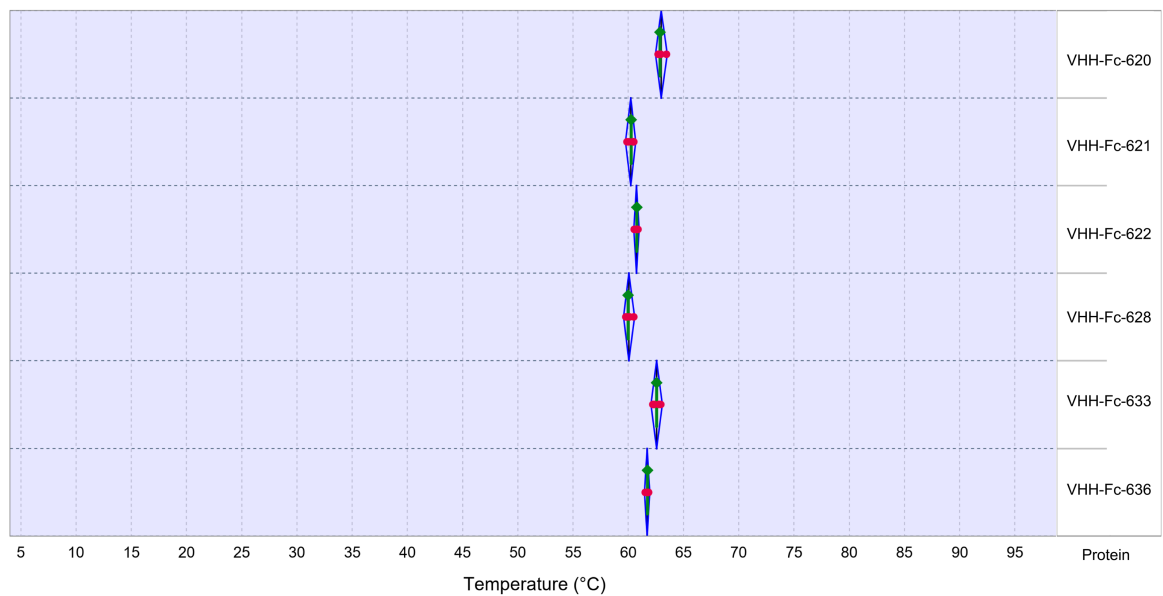

**Supplementary Figure S1. Biochemical characterization of VHH-Fcs**

**(a)** Analytical size-exclusion chromatography (SEC) traces of purified VHH-Fcs. SDS-PAGE analysis (10%) under reducing (R) and non-reducing (NR) conditions confirms the integrity of the VHH-Fcs.

**(b)** Thermal stability of VHH-Fc fusion proteins determined by differential scanning fluorimetry. Melting temperatures ($T_m$) of six VHH-Fc constructs (VHH-Fc 620, 621, 622, 628, 633, and 636) were determined using the Protein Thermal Shift™ assay ($n = 4$ technical replicates per protein). Red dots represent the $T_m$ values of individual replicates, the green diamond indicates the median $T_m$, and the blue whiskers indicate the full range of $T_m$ values across replicates.

### Supplementary Figure S2

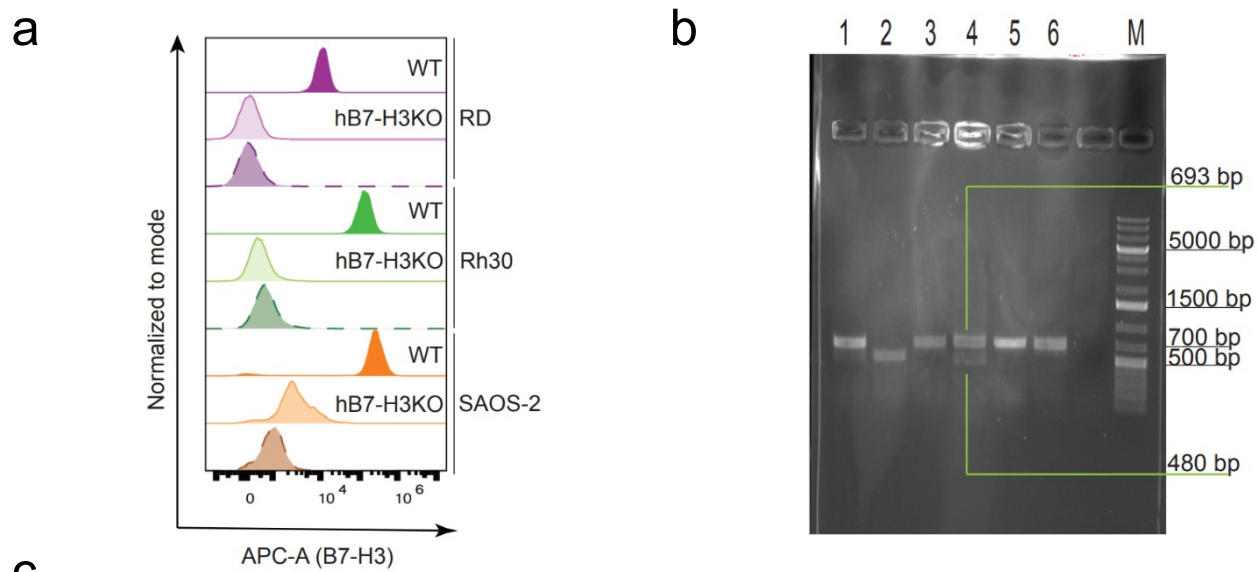

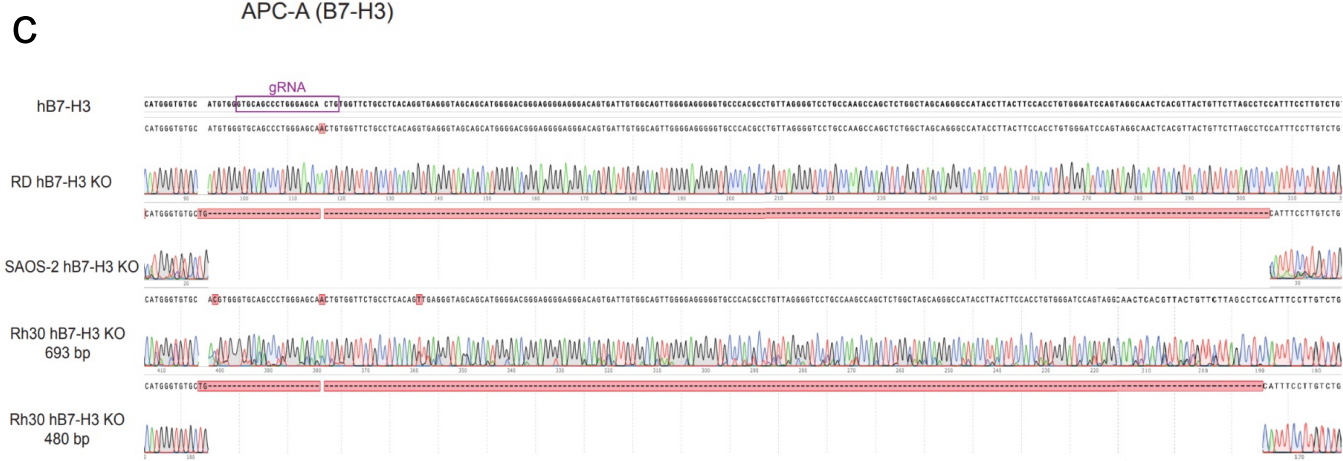

#### **Supplementary Figure S2. Validation of B7-H3 knockout in Saos-2, RH30, and RD cell lines**

**(a)** Flow cytometric analysis of B7-H3 surface expression in wild-type (WT) and CRISPR/Cas9-generated B7-H3 knockout (KO) single-cell clones across RD, RH30, and Saos-2 cell lines. Cells were stained with an anti-B7-H3 antibody and histograms are normalized to mode. Dashed lines represent isotype control staining. Loss of detectable B7-H3 surface expression confirms successful KO in all three cell lines.

**(b)** Genomic PCR validation of B7-H3 KO clones. Genomic DNA was amplified using primers flanking the gRNA target site. In RD and RH30 KO clones, a band at ~693 bp, identical in size to the WT band, was retained due to a small insertion at the gRNA cut site. Additionally, RH30 KO clones harbor a second allele carrying a large deletion, producing a smaller ~480 bp band. In Saos-2 KO clones, only the ~480 bp band is observed, reflecting a large deletion on both alleles. Together, these banding patterns indicate biallelic disruption of the B7-H3 locus in all three KO clones. Lane assignments: 1. Saos-2 WT, 2. Saos-2 B7-H3 KO, 3. RH30 WT, 4. RH30 B7-H3 KO, 5. RD WT, 6. RD B7-H3 KO; M: GeneRuler 1 kb Plus DNA ladder.

**(c)** Sanger sequencing chromatograms of the PCR bands isolated in (b). The reference sequence is shown at the top with the gRNA target site highlighted in purple. In RD and RH30 KO clones, sequencing of the 693 bp band reveals a small insertion at the gRNA cut site, while the 480 bp band from RH30 and Saos-2 KO clones shows a large deletion spanning the target site (red dashed region). Together, these results confirm biallelic frameshift mutations at the B7-H3 locus in all three knockout clones.

Supplementary Figure S3

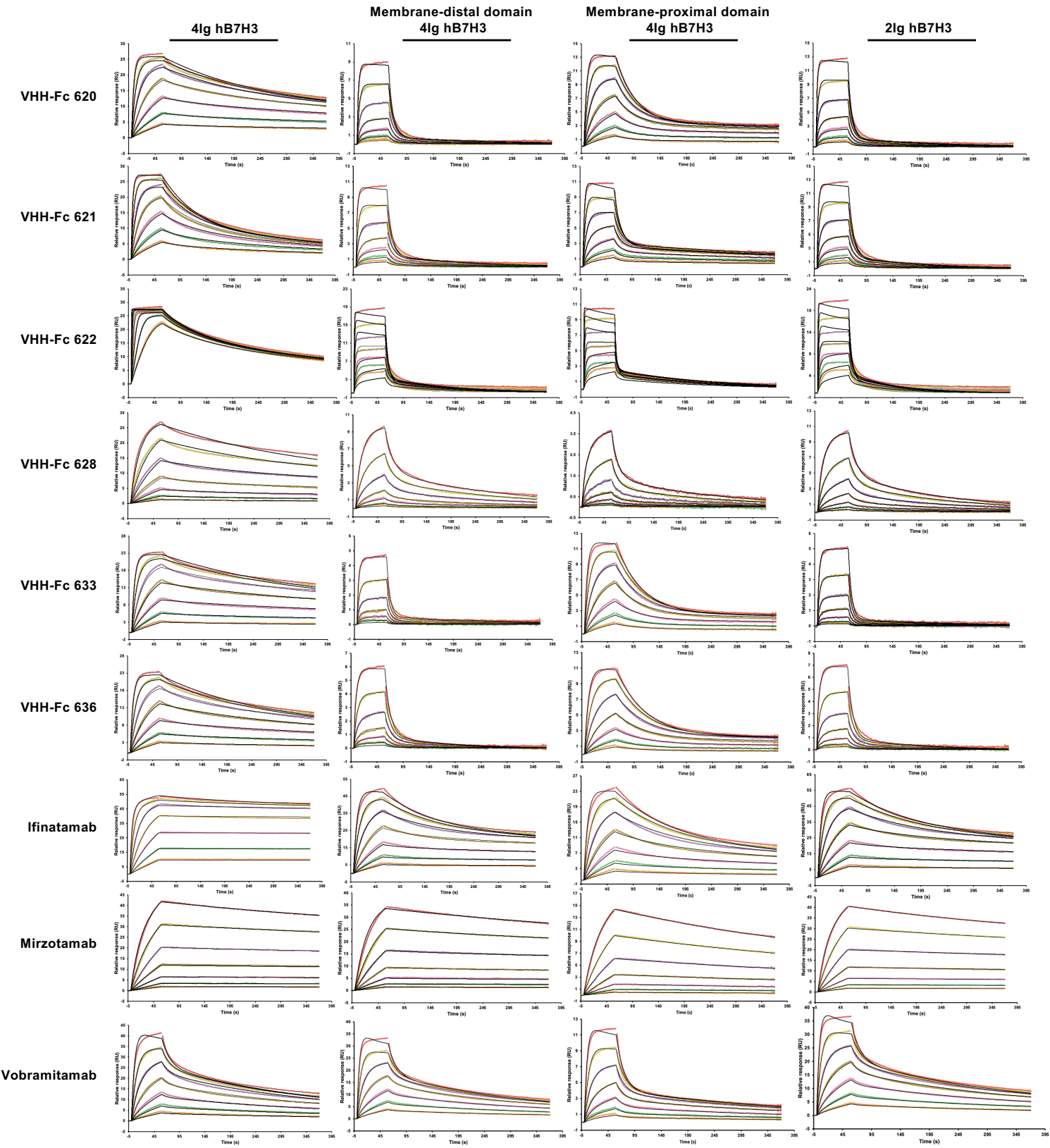

**Supplementary Figure S3. SPR-based binding affinity analysis of VHH-Fcs and benchmarking mAbs**

Sensorgrams of VHH-Fc 620, 621, 622, 628, 633, 636, Ifinatamab, Mirzotamab, Vobramitamab binding to 4Ig hB7-H3, membrane-distal and membrane-proximal domains of 4Ig hB7-H3 and 2Ig hB7-H3. Tested mAbs concentrations ranged from 100 to 1.56 nM in a twofold serial dilution (100, 50, 25, 12.5, 6.25, 3.125, and 1.56 nM), corresponding to the following sensorgram colors: salmon, yellow, purple, brown, pink, green, and orange, respectively. The sensorgrams shown are representative of three independent experiments with similar results.

Supplementary Figure S4

VHH-Fc 620

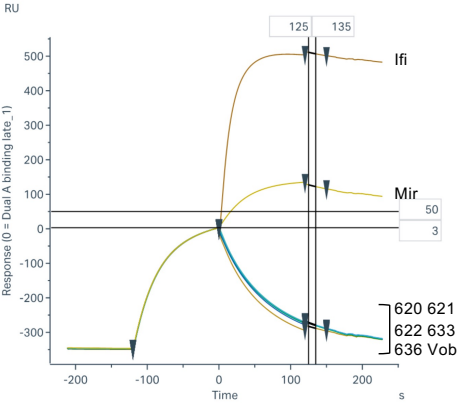

VHH-Fc 621

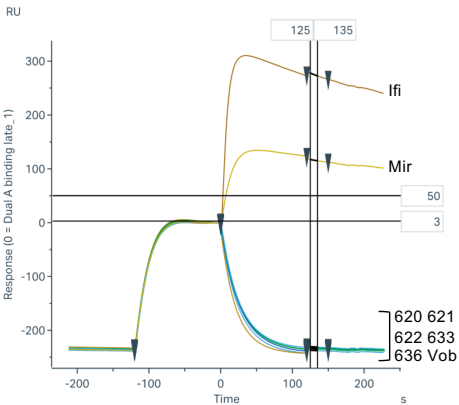

VHH-Fc 622

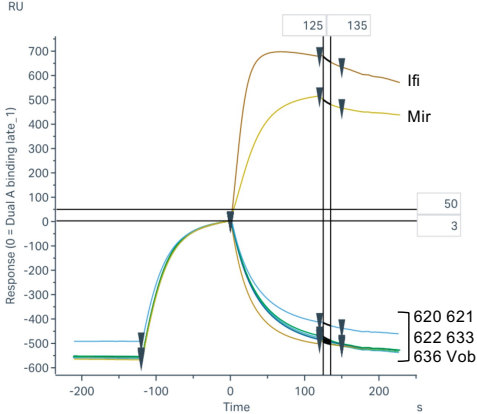

VHH-Fc 633

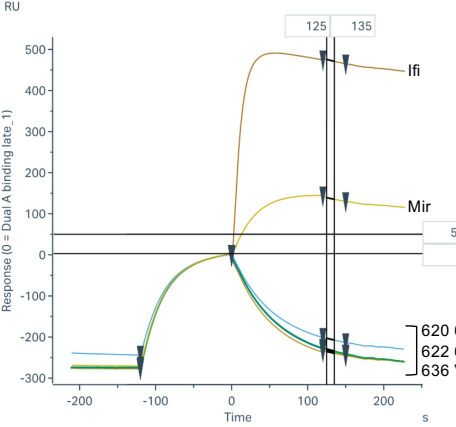

VHH-Fc 636

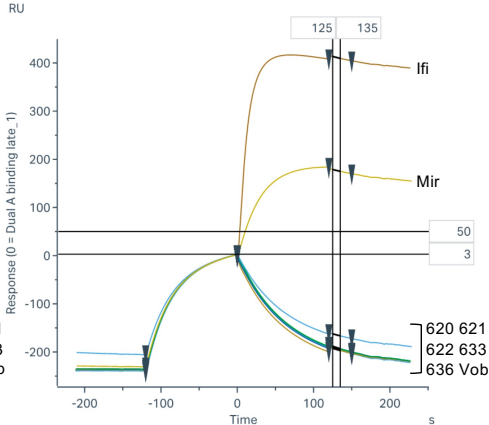

Vobramitamab

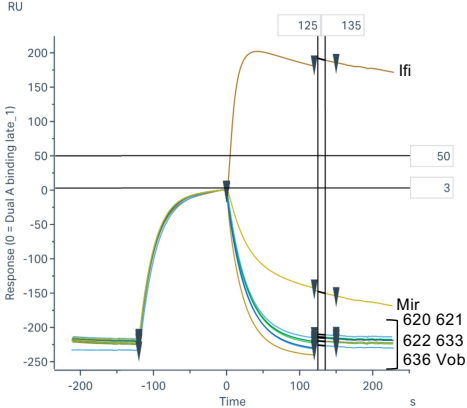

Mirzotamab

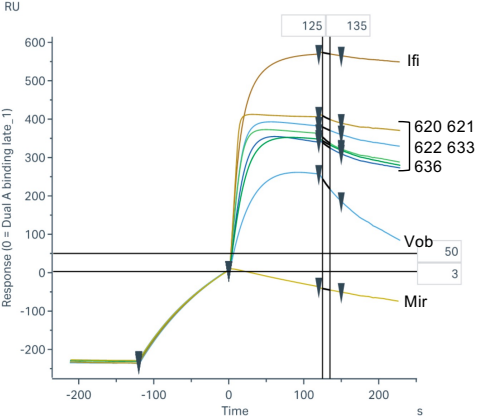

Ifinatamab

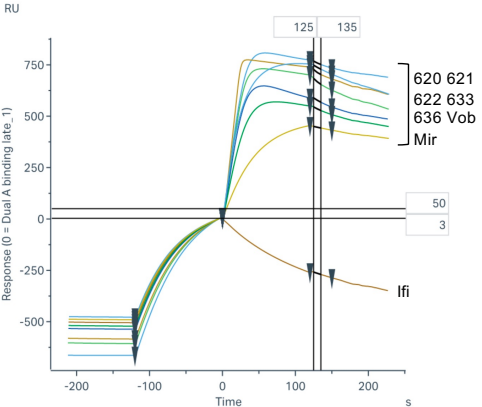

**Supplementary Figure S4. SPR-based epitope binning of VHH-Fcs and benchmarking mAbs**

SPR sensorgrams of all binder combinations analyzed as either capture binder (indicated in panel title) or secondary binder to determine epitope bins. VHH-Fc 628 is excluded, as it bound the membrane-proximal domain too weakly for reliable binning (see Supplementary Figure S5). Sensorgrams show injections of the membrane-proximal domain of 4Ig hB7-H3 (starting at -120 s), followed by injection of the second binder (starting at ~0 s). A response above the upper threshold line indicates simultaneous binding, whereas a response at or below baseline indicates competition. Vob, vobramitamab; Mir, mirzotamab; Ifi, ifinatamab.

Supplementary Figure S5

VHH-Fc 620

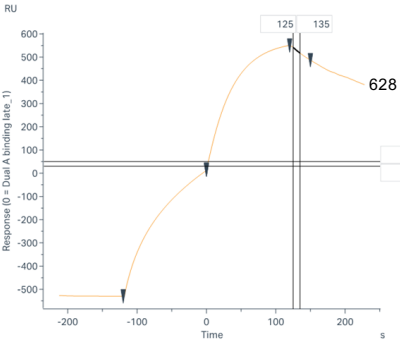

VHH-Fc 621

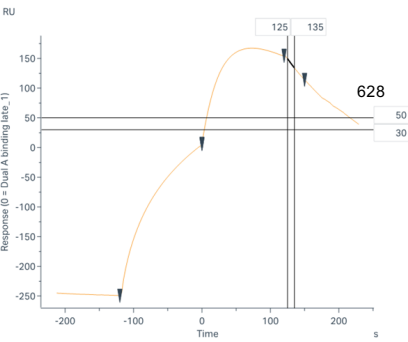

VHH-Fc 622

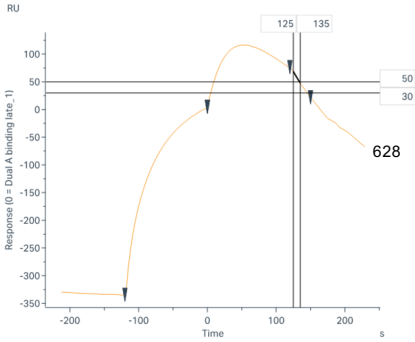

VHH-Fc 628

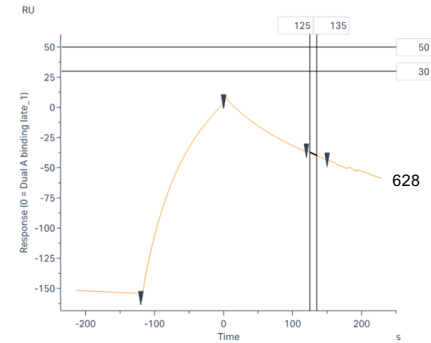

VHH-Fc 633

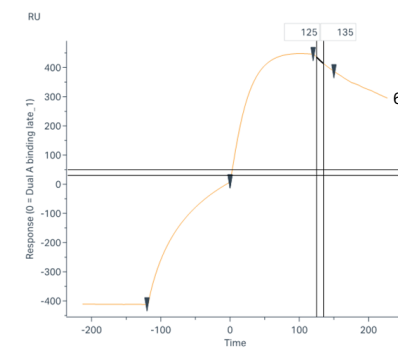

VHH-Fc 636

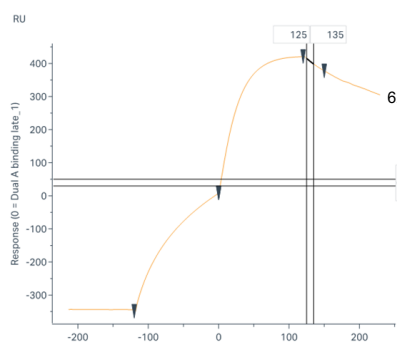

Vobramitamab

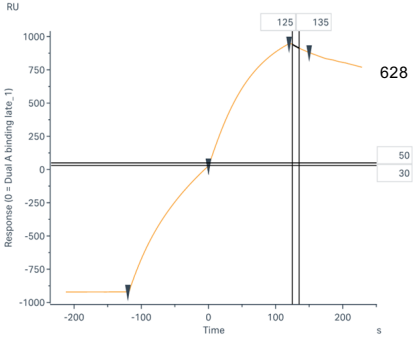

Mirzotamab

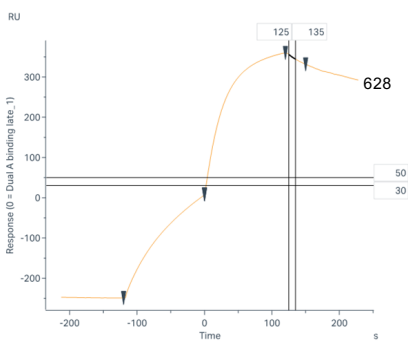

Ifinatamab

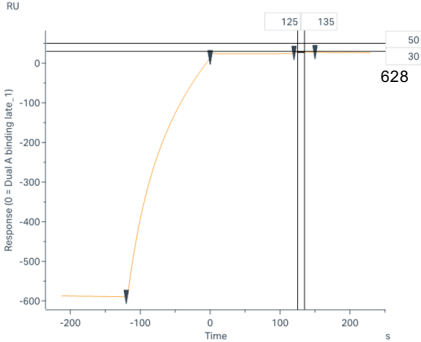

**Supplementary Figure S5. SPR-based epitope binning of VHH-Fc 628**

SPR sensorgrams of all combinations of VHH-Fc 628 with the VHH-Fc panel and benchmarking mAbs, analyzed as either capture binder (indicated in the panel title) or secondary binder. Sensorgrams show injection of full-length 4Ig hB7-H3 (starting at -120 s), followed by injection of the second binder (starting at ~0 s). Binning for VHH-Fc 628 was performed on full-length 4Ig hB7-H3 rather than the isolated membrane-proximal domain, as VHH-Fc 628 bound the isolated domain too weakly for reliable binning.

Supplementary Figure S6

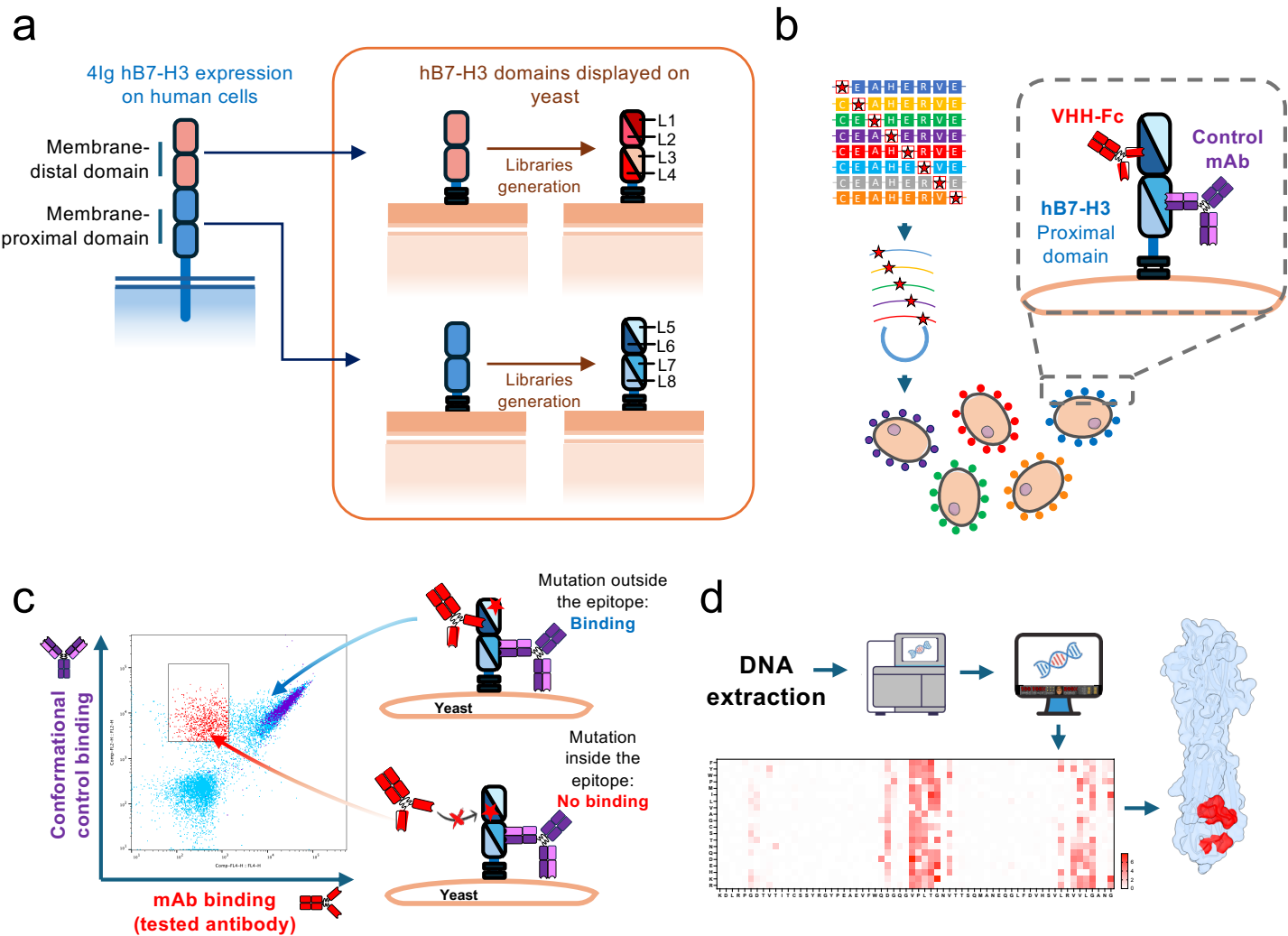

**Supplementary Figure S6. Concept of epitope mapping by deep mutational scanning, yeast surface display and next generation sequencing**

**(a)** Schematic of 4Ig hB7-H3 at the mammalian cell surface and of its two IgV-IgC domain pairs displaying separately on yeast, so that each binder could engage only a single copy of its epitope. For the membrane-distal domain (amino acids 29–238) and the membrane-proximal domain (amino acids 243–456), numbered according to UniProt Q5ZPR3, we constructed four DNA libraries each, containing clones with a single mutation introduced by an NNK codon at each position (designated L1–L4 for the membrane-distal domain and L5–L8 for the membrane-proximal domain).

**(b)** The libraries were transformed into *S. cerevisiae* cells using gap repair recombination.

**(c)** Following induction, each library was incubated with the binder under investigation, together with a conformational control antibody. During FACS analysis, clones that lost recognition by the binder under study while retaining conformational control antibody staining (red gates) were sorted.

**(d)** DNA from the sorted population was sequenced using Illumina next-generation sequencing (NGS). Bioinformatic analysis was used to generate heat maps of enrichment, defined as the fold change in amino acid substitution frequency between sorted and unsorted populations. Amino acid residues identified as critical were then mapped in color onto the AlphaFold 3 prediction of hB7-H3, defining the functional epitope.

Supplementary Figure S7

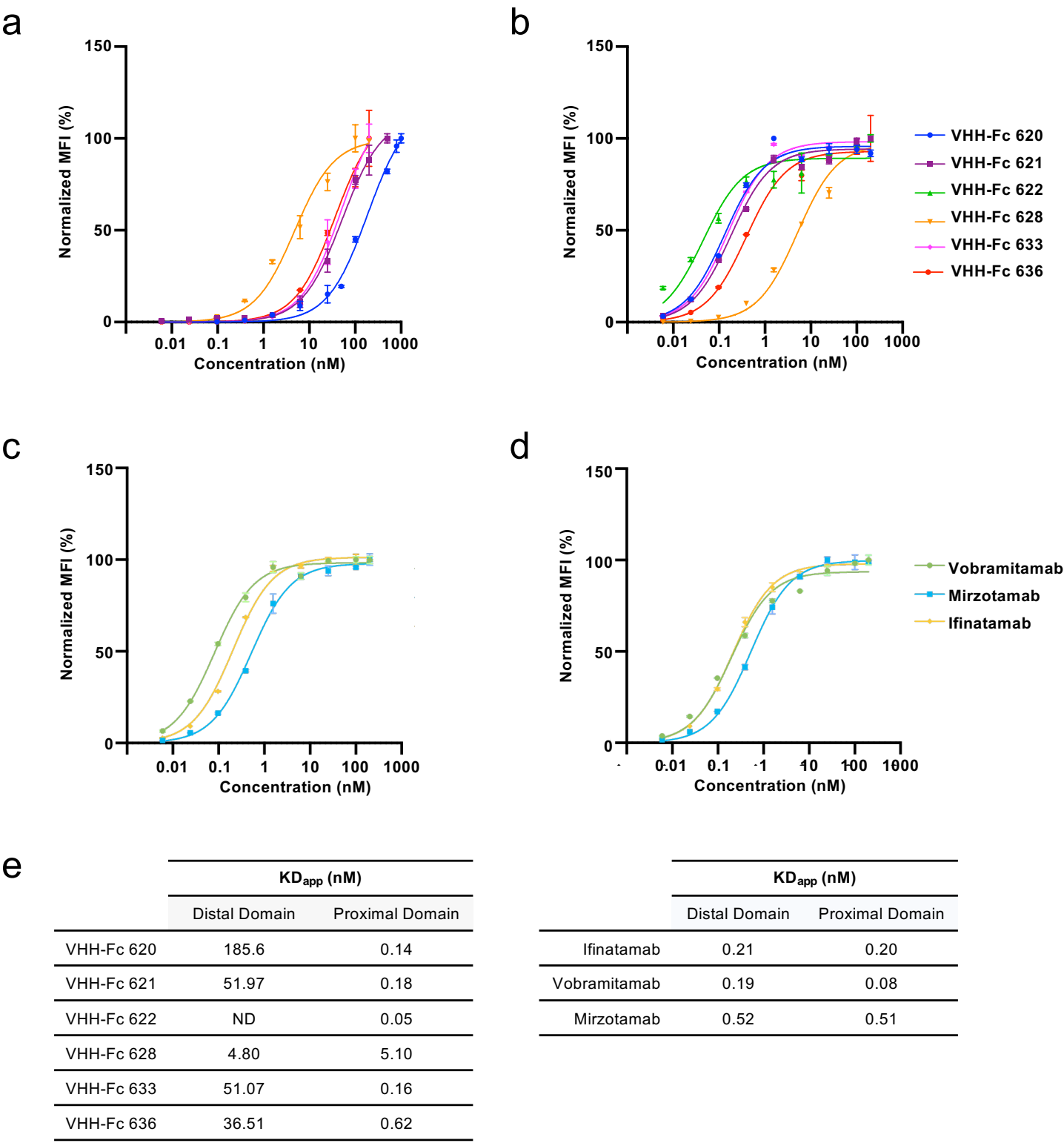

**Figure S7. Apparent affinities of the VHH-Fcs and benchmarking mAbs for the membrane-proximal and membrane-distal domains of 4Ig hB7-H3 displayed on yeast**

**(a)** Binding curves of the VHH-Fcs to yeast displaying the hB7-H3 membrane-distal domain, measured by flow cytometry.

**(b)** Binding curves of the VHH-Fcs to yeast displaying the hB7-H3 membrane-proximal domain.

**(c)** Binding curves of the benchmarking mAbs to the membrane-distal domain.

**(d)** Binding curves of the benchmarking mAbs to the membrane-proximal domain.

**(e)** Apparent binding affinities ($KD_{app}$) values calculated from the curves in (a–d) by non-linear regression in GraphPad Prism 10.

Means ± SD of duplicate values (technical replicates) are shown. MFI, mean fluorescence intensity; ND, not determined.

Supplementary Figure S8

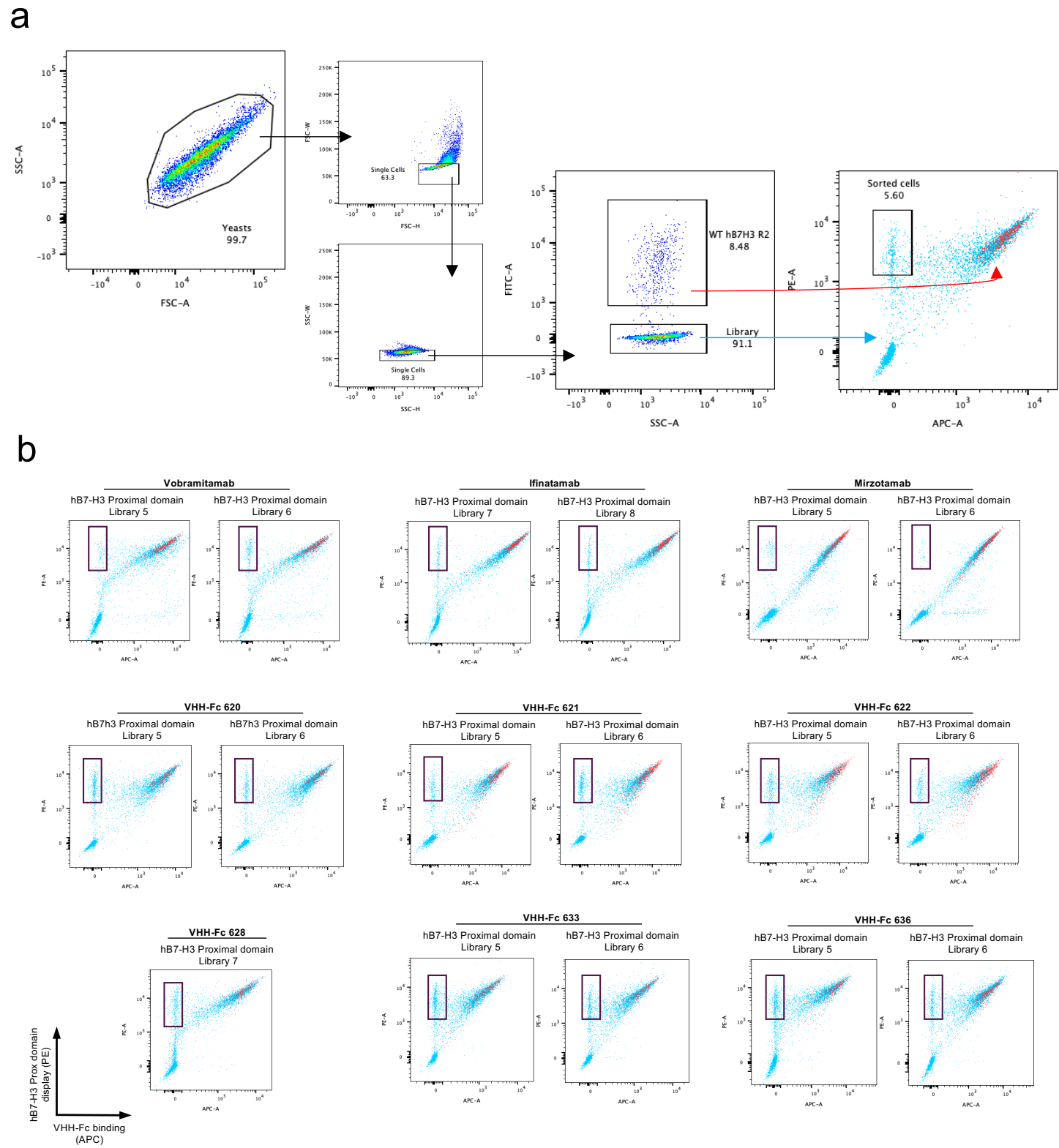

**Supplementary Figure S8. Flow cytometry gating strategy and sorting of yeast populations**

**(a)** Cells were initially gated based on forward scatter (FSC-A) and side scatter (SSC-A) to isolate the yeast population. Doublets were excluded by sequential gating using FSC-H versus FSC-W and SSC-H versus SSC-W to obtain a single-cell population. Single cells were then analyzed by fluorescence (FITC-A vs SSC-A), enabling separation of the library population (blue) from the wild-type (WT) hB7-H3 membrane-proximal domain population (red). Cell populations were further analyzed based on APC-A (VHH-Fc/mAb binding) versus PE-A (hB7-H3 display), as shown in the final dot plot. The sorted subpopulation (black box) corresponds to cells exhibiting PE signal in the absence of APC signal.

**(b)** Dot plots of yeast displaying the hB7-H3 membrane-proximal domain stained with the VHH-Fc and benchmarking mAbs. The sorted cell population for each binder is indicated by the black box. Libraries used for each binder are indicated above each plot.

Supplementary Figure S9

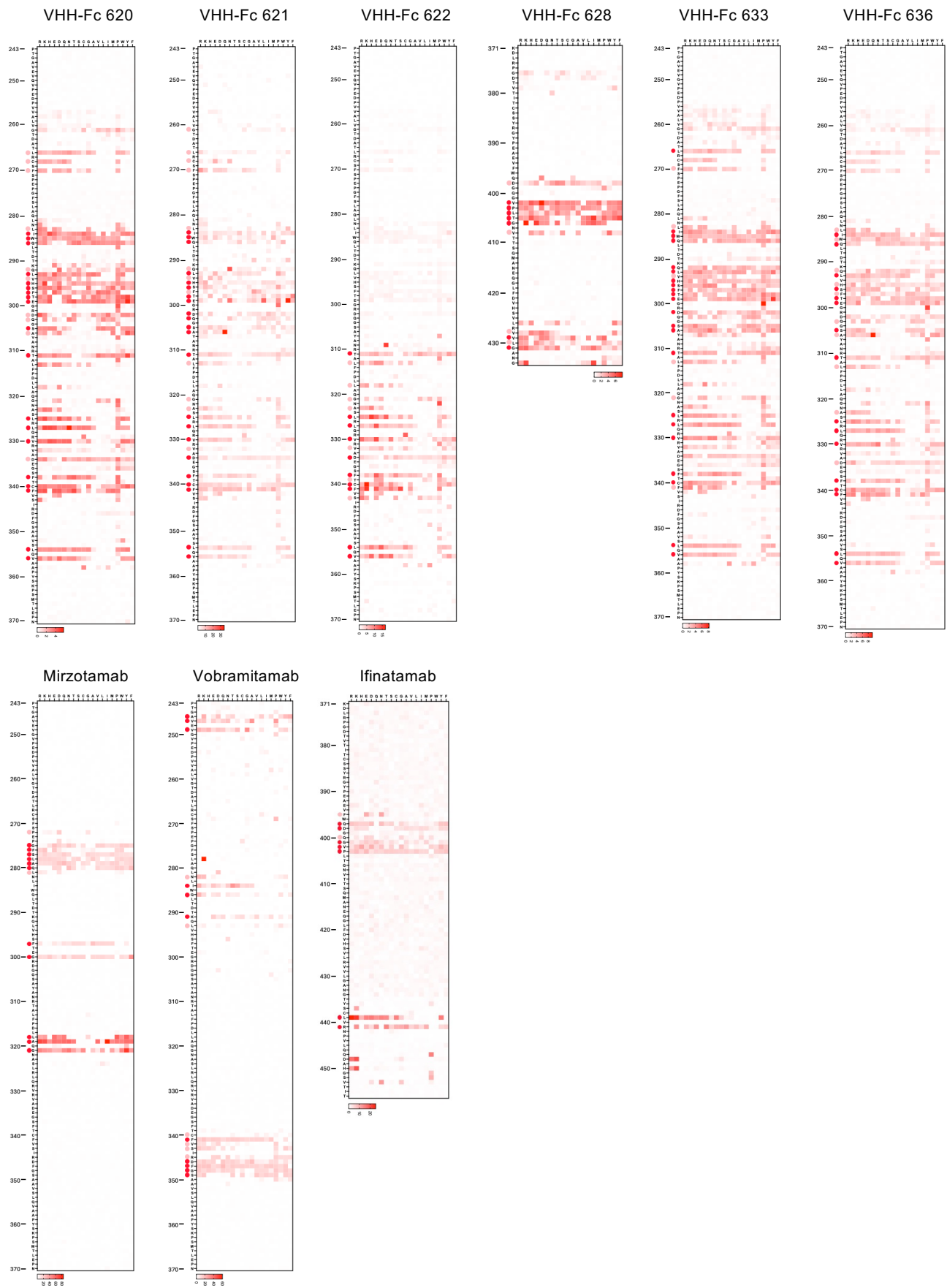

**Supplementary Figure S9. Heatmaps depicting the functional epitopes of hB7-H3 binders on the membrane-proximal domain of hB7-H3**

Heatmap of the hB7-H3 membrane-proximal domain (amino acids 243-456, Q5ZPR3 · CD276\_HUMAN ) showing the effect of amino acid substitution on VHH-Fc and mAb binding. Dark red marks residues where $\geq 50\%$ of the substitutions tested (n = 19) disrupted binding, while light red marks residues where $\geq 25\%$ of substitutions disrupted binding.

Supplementary Figure S10

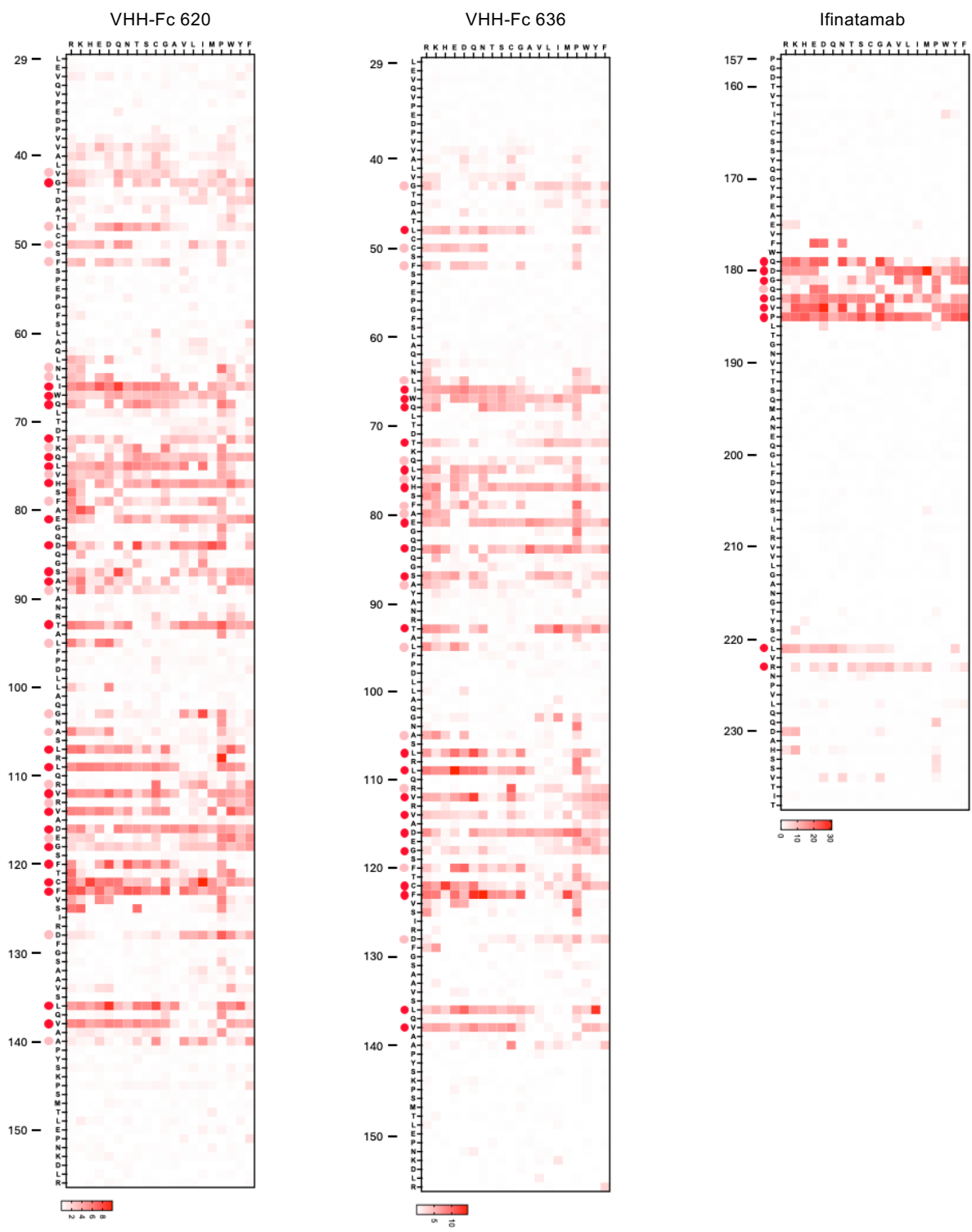

**Supplementary Figure S10. Heatmaps depicting the functional epitopes of hB7-H3 binders on the membrane-distal domain of hB7-H3**

Heatmap of the hB7-H3 membrane-distal domain (amino acids 29-238, Q5ZPR3 · CD276\_HUMAN ) showing the effect of amino acid substitution on VHH-Fc and mAb binding. Dark red marks residues where $\geq 50\%$ of the substitutions tested (n = 19) disrupted binding, while light red marks residues where $\geq 25\%$ of substitutions disrupted binding.

Supplementary Figure S11

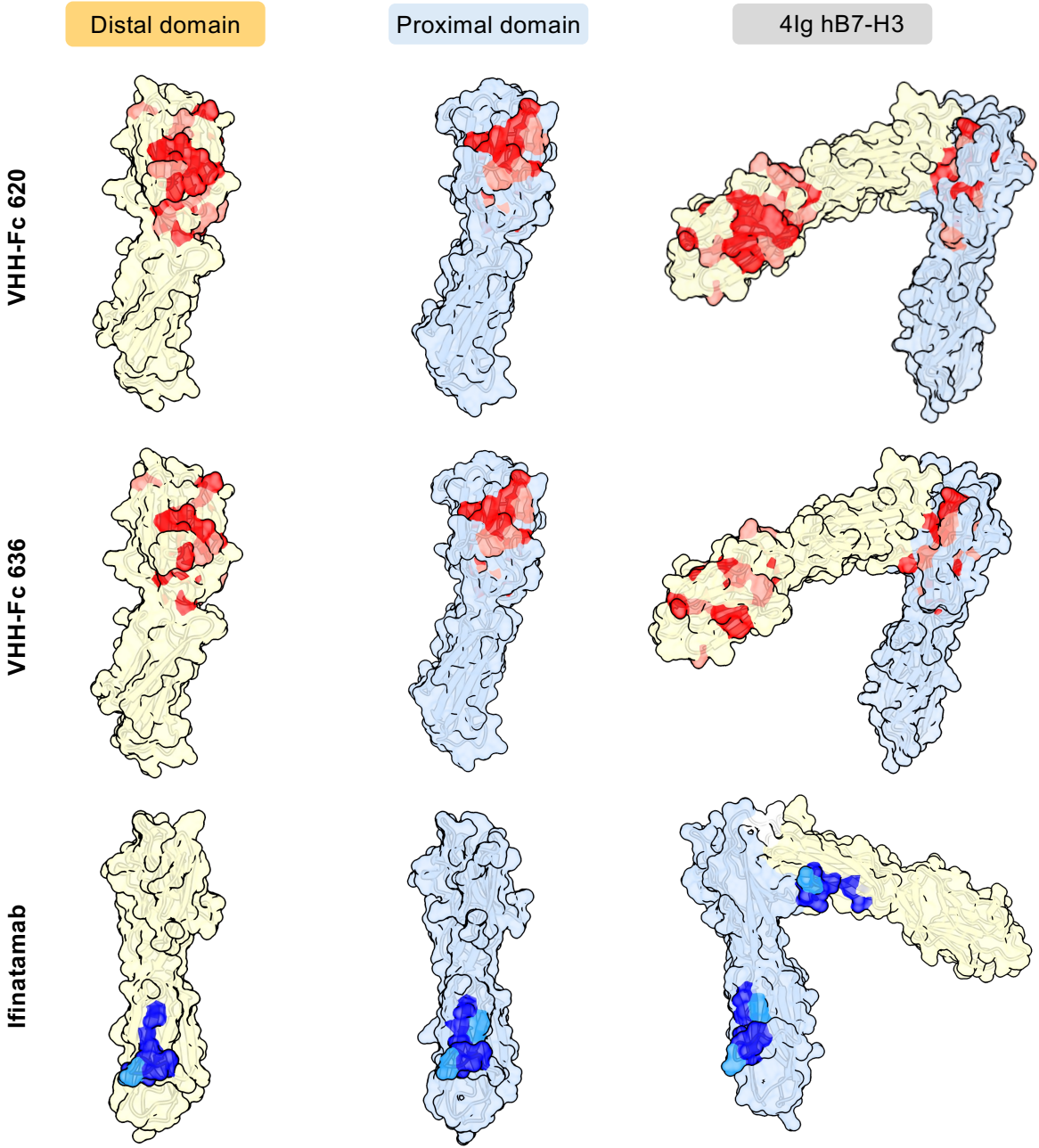

**Supplementary Figure S11. Functional epitopes of hB7-H3 binders mapped onto the AlphaFold3 structures of the membrane-proximal domain, membrane-distal domain, and 4Ig hB7-H3**

Surface representations of the membrane-proximal and membrane-distal domains of hB7-H3, as well as the full-length 4Ig hB7-H3 model predicted by AlphaFold3, highlighting putative amino acid residues involved in binding to B7-H3–specific VHH-Fc 620, VHH-Fc 636, and ifinatamab. Residues are color-coded according to the impact of mutagenesis on VHH-Fc/mAb binding: dark colors (red or blue) indicate positions where $\geq 50\%$ of substitutions ($n = 19$) disrupt binding, whereas lighter colors (red or blue) indicate positions where $\geq 25\%$ of substitutions disrupt binding.

Supplementary Figure S12

a

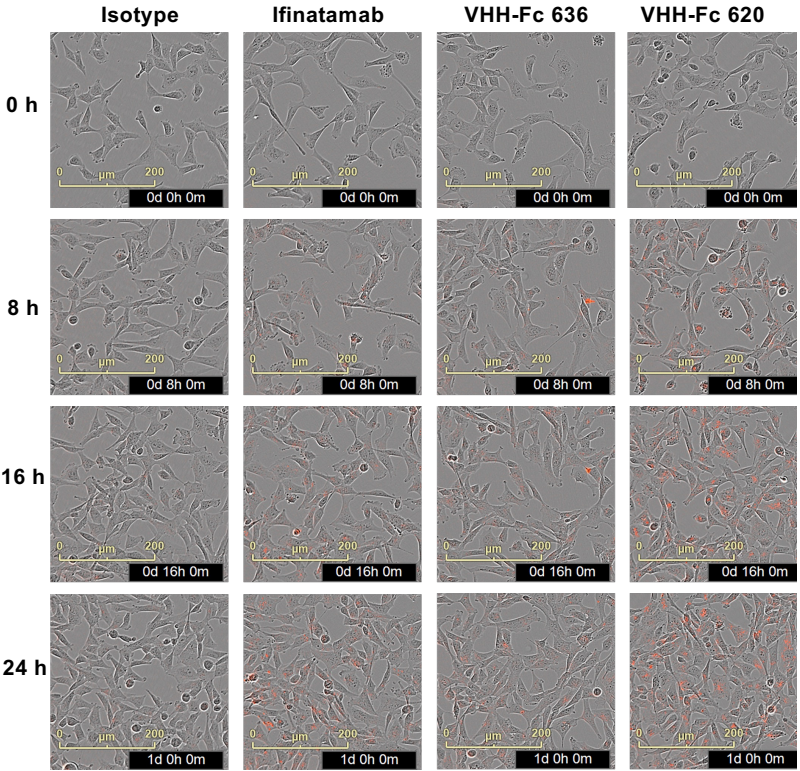

b

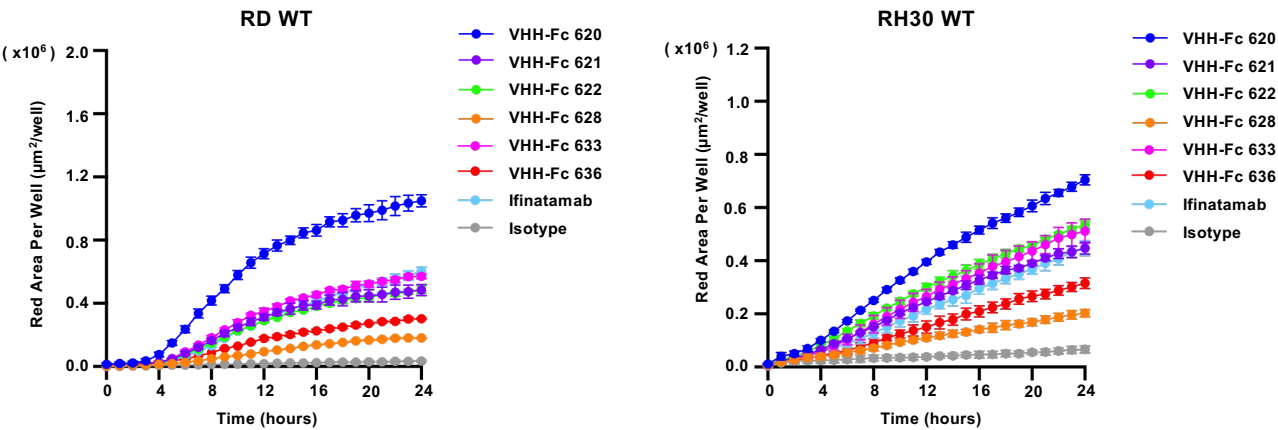

c

**Supplementary Figure S12. Real-time analysis of VHH-Fc internalization using the IncuCyte live-cell imaging system**

**(a)** Representative Incucyte images (phase contrast with red fluorescence overlay) of RD WT cells at 0, 8, 16, and 24 h after treatment with 13.5 nM isotype control, ifinatamab, VHH-Fc 636, or VHH-Fc 620. Red fluorescence indicates intracellular accumulation of labeled binders, reflecting internalization over time.

**(b)** Quantitative kinetic analysis of VHH-Fc and ifinatamab internalization at 6.75 nM in RD WT and RH30 WT cells, measured as red object area per well over 24 h using the Incucyte system. Data are shown as mean ± SD from one representative assay of three independent experiments with similar results, each including three technical replicates.

**(c)** Endpoint quantification at 24 h of the data shown in (b), normalized to ifinatamab. Bars represent mean ± SD from three independent experiments, each including three technical replicates. Statistical significance was determined by one-way ANOVA with Dunnett's multiple comparisons test against ifinatumab (\* $p < 0.05$, \*\* $p < 0.01$, \*\*\* $p < 0.001$, \*\*\*\* $p < 0.0001$; ns, not significant) in GraphPad Prism 10.

Supplementary Figure S13

a

VHH-Fc 620\_MC\_GGFG\_PAB\_Exatecan

|  | VHH-Fc 620 |  |  | VHH-Fc 620_MC_GGFG_PAB_Exatecan |  |  |
| --- | --- | --- | --- | --- | --- | --- |
|  | Mass | Intensity | Intensity% | Mass | Intensity | Intensity% |
| H0 | 40674.08483 | 9.54E+09 | 46.6% | 40678.8281 | 8.28E+08 | 13.8% |
|  | 41603.53915 | 1.10E+10 | 53.4% | 41608.38864 | 1.49E+09 | 24.9% |
| H1 |  |  | 0.0% | 41775.37283 | 7.10E+08 | 11.8% |
|  |  |  | 0.0% | 42704.22259 | 8.56E+08 | 14.3% |
| H2 |  |  | 0.0% | 42871.48773 | 8.42E+08 | 14.0% |
|  |  |  | 0.0% | 43800.45372 | 1.28E+09 | 21.2% |
| H3 |  |  | 0.0% |  |  | 0.0% |
|  |  |  | 0.0% |  |  | 0.0% |
| DAR |  |  | 0.00 |  |  | 1.9 |

| Retention time (min) | Area (mAU*s) | Relative area (%) | Purity (%) |
| --- | --- | --- | --- |
| 8.400 | 164661 | 10.399 | 89.60 |
| 8.935 | 1418821 | 89.601 |  |

B

Ifinatamab\_MC\_GGFG\_PAB\_Exatecan

|  | Ifinatamab |  |  | Ifinatamab_MC_GGFG_PAB_Exatecan |  |  |
| --- | --- | --- | --- | --- | --- | --- |
|  | Mass | Intensity | Intensity% | Mass | Intensity | Intensity% |
| L0 | 23313.9766 | 7.65E+06 | 64.1% | 23316.9805 | 1.30E+07 | 36.1% |
|  | 23317.9199 | 4.29E+06 | 35.9% | 23320.9102 | 9.05E+06 | 25.1% |
| L1 |  |  |  | 24412.9434 | 8.00E+06 | 22.2% |
|  |  |  |  | 24415.793 | 5.98E+06 | 16.6% |
| H0 | 51250.8633 | 3.46E+07 | 66.5% | 51109.6328 | 1.84E+06 | 14.5% |
|  | 51104.6875 | 1.74E+07 | 33.5% | 51255.4141 | 3.47E+06 | 27.4% |
| H1 |  |  |  | 52205.6016 | 1.24E+06 | 9.8% |
|  |  |  |  | 52351.4297 | 4.16E+06 | 32.8% |
| H2 |  |  |  | 52205.6016 | 1.24E+06 | 9.8% |
|  |  |  |  | 52351.4297 | 4.16E+06 | 32.8% |
| DAR |  |  | 0.00 |  |  | 2.25 |

| Retention time (min) | Area (mAU*s) | Relative area (%) | Purity (%) |
| --- | --- | --- | --- |
| 6.757 | 61.18 | 1.82 | 98.18 |
| 7.510 | 330.85 | 98.18 |  |

**Supplementary Figure S13. Mass spectrometry and HPLC characterization of VHH-Fc 620 and ifinatamab conjugated with exatecan**

**(a)** Characterization of VHH-Fc 620–MC-GGFG-PAB-exatecan. Deconvoluted intact mass spectrum (top left) and corresponding mass/intensity summary table (top right) of unconjugated VHH-Fc 620 and the conjugate, showing the distribution of drug-loaded species (H0–H3) and the calculated average drug-to-antibody ratio (DAR = 1.9). HPLC chromatogram (bottom left) and retention time, peak area and purity table (bottom right) of the conjugate, indicating 89.60% purity.

**(b)** Characterization of ifinatamab–MC-GGFG-PAB-exatecan. Deconvoluted mass spectrum under reducing conditions (top left) and summary table (top right) of unconjugated ifinatamab and the conjugate, showing conjugated light chain (L0/L1) and heavy chain (H0–H2) species and the calculated DAR (2.25, light and heavy chain combined). HPLC chromatogram (bottom left) and retention time, peak area and purity table (bottom right) of the conjugate, indicating 98.18% purity.

Conjugation and analysis were performed by MedChemExpress.

Supplementary Figure S14

**b**

| MFI FL3 |  |  |  |  |
| --- | --- | --- | --- | --- |
|  | RD | RH30 | MG63 | Saos-2 |
| KO | 331 ± 36 | 359 ± 32 | 335 ± 4 | 323 ± 27 |
| WT | 33106 ± 4016 | 57833 ± 6405 | 31591±283 | 18416 ± 1823 |
| OV | 423427 ± 16580 | 203431± 4881 | 276031±14103 | 271114 ± 9075 |

**Supplementary Figure S14. Flow cytometric characterization of B7-H3 surface expression in knockout, wild-type, and overexpression cell lines**

**(a)** Representative flow cytometry histograms showing cell-surface B7-H3 expression in RD, RH30, MG63, and Saos-2 cells. B7-H3 knockout (KO), wild-type (WT), and B7-H3-overexpressing (OV) cells are shown in grey, cyan, and red, respectively. The X-axis represents B7-H3-associated fluorescence intensity and the Y-axis the normalized cell count. Cells were stained with first antibody ifiantamab and second antibody donkey anti-Human IgG (H+L) Cross-Adsorbed Secondary Antibody, DyLight™ 650 (Thermo Fisher, SA5-10129). The KO clones are those validated in Supplementary Figure S2. Representative results from one of three independent experiments are shown.

**(b)** Mean fluorescence intensity (MFI) of B7-H3-associated fluorescence for KO, WT, and OV cells across the four cell lines. Values are mean ± SD of three independent experiments.

Supplementary Figure S15

**b**

| IC50 (nM) |  |  |  |
| --- | --- | --- | --- |
| Cell line | 620-Exatecan | Ifinatomab-Exatecan | Exatecan |
| RH30 KO | N | N | 0.31 (95 % CI :0.19-0.69) |
| RH30 WT | N | N | 0.38 (95 % CI :0.32-0.45) |
| RH30 OV | 19.39 (95 % CI :14.31-28.58) | 0.68 (95 % CI :0.43-1.00) | 0.42 (95 % CI :0.25-0.60) |
| RD KO | N | N | 0.63 (95 % CI :0.49-0.84) |
| RD WT | N | N | 0.64 (95 % CI :0.55-0.78) |
| RD OV | 6.71 (95 % CI :5.92-7.87) | 0.35 (95 % CI :0.31-0.40) | 0.70 (95 % CI :0.60-0.81) |
| MG63 KO | N | N | 0.59 (95 % CI :0.52-0.67) |
| MG63 WT | N | N | 0.56 (95 % CI :0.47-0.6) |
| MG63 OV | 3.03 (95 % CI :2.45-3.82) | 0.88 (95 % CI :0.79-1) | 0.46 (95 % CI :0.39-0.52) |
| Saos-2 KO | N | N | 0.81 (95 % CI :0.76-0.92) |
| Saos-2 WT | N | N | 1.13 (95 % CI :1.01-1.28) |
| Saos-2 OV | 31.27 (95 % CI :24.14-45.32) | 3.36 (95 % CI :2.45-5.44) | 1.1 (95 % CI :0.96-1.26) |

**Supplementary Figure S15. Exatecan conjugation preserves B7-H3 binding affinity and mediates B7-H3-dependent cytotoxicity in sarcoma cell lines**

**(a)** Concentration–response curves showing the effects of VHH-Fc 620–exatecan, ifinatamab–exatecan, the corresponding unconjugated binders, and free exatecan on cell viability in MG63 and Saos-2 cells, each as wild-type (WT), B7-H3-knockout (KO), and B7-H3-overexpressing (OV) derivatives. Cell viability was measured after 120 h using CellTiter-Glo® 2.0 and normalized to that of untreated controls. Data are presented as mean ± SD from three independent experiments, each performed in technical triplicate. Concentration–response curves were fitted by nonlinear regression using the variable-slope four-parameter logistic model ([inhibitor] vs. response) in GraphPad Prism 10.

**(b)** Summary of IC<sub>50</sub> values and corresponding 95% confidence intervals derived from the nonlinear regression analysis in (a) and in Figure 5c. "N" indicates that the IC<sub>50</sub> could not be reliably determined because a complete concentration–response curve was not achieved within the tested concentration range up to 200 nM.
